# Bringing up-to-date the toolkit for the catabolism of aromatic compounds in fungi: the unexpected 1,2,3,5–tetrahydroxybenzene central pathway

**DOI:** 10.1101/2023.06.06.543907

**Authors:** Tiago M. Martins, Artur Bento, Celso Martins, Ana S. Tomé, Carlos J.S. Moreira, Cristina Silva Pereira

**Affiliations:** Instituto de Tecnologia Química e Biológica António Xavier, Universidade Nova de Lisboa (ITQB NOVA), Av. da República, 2780-157 Oeiras, Portugal

**Keywords:** Gallate, 1,2,3,5–Tetrahydroxybenzene, RNA-seq, Aromatic compounds catabolism, *Aspergillus terreus*, *Aspergillus nidulans*, Saprophytic Ascomycota fungi

## Abstract

Saprophytic fungi are able to catabolize many plant-derived aromatics, including, for example, gallate. The catabolism of gallate in fungi is assumed to depend on the five main central pathways, *i.e.*, of the central intermediates’ catechol, protocatechuate, hydroxyquinol, homogentisate, and gentisate, but a definitive demonstration is lacking. To shed light on this process, we analyzed the transcriptional reprograming of the growth of *Aspergillus terreus* on gallate compared with acetate as the control condition. Surprisingly, the results revealed that the five main central pathways did not exhibit significant positive regulation. Instead, an in-depth analysis identified four highly expressed and upregulated genes that are part of a conserved gene cluster found in numerous species of fungi, though not in *Aspergilli*. The cluster comprises a monooxygenase gene and a fumarylacetoacetate hydrolase-*like* gene, which are recognized as key components of catabolic pathways responsible for aromatic compound degradation. The other two genes encode proteins with no reported enzymatic activities. Through functional analyses of gene deletion mutants, the conserved short protein with no known domains could be linked to the conversion of the novel metabolite 5-hydroxydienelatone, whereas the DUF3500 gene likely encodes a ring-cleavage enzyme for 1,2,3,5–tetrahydroxybenzene. These significant findings establish the existence of a new 1,2,3,5-tetrahydroxybenzene central pathway for the catabolism of gallate and related compounds (*e.g.,* 2,4,6-trihydroxybenzoate) in numerous fungi where this catabolic gene cluster was observed.

**IMPORTANCE:** The lignin found in various economically significant plants, such as major grains like rice, wheat, and maize, comprises a substantial portion of syringyl units (up to 60%). As a result, the future utilization of residues from these plants in biorefineries will yield significant quantities of syringyl derivatives. However, our understanding of how fungi degrade these syringyl derivatives is to date scarce and mostly relies on unproven assumptions. Our study, demonstrates the existence of a new 1,2,3,5-tetrahydroxybenzene central intermediate for the catabolism of gallate in numerous fungi. This finding expands the toolkit of central pathways, proving that the generalized assumption that gallate catabolism depends on the previously known five main central pathways was incorrect. This research reveals a novel crucial central pathway of great ecological and biotechnological importance, not only for fungi but also potentially for bacteria.

## Introduction

Saprophytic fungi play an invaluable role in the turnover of the elements, including those present in highly abundant plant phenolic sources such as lignin and tannins (1). Understanding the catabolism of aromatics, for example, gallate, a major intermediate of the metabolism of lignin constituents, esterified polysaccharides, and hydrolyzable tannins (1), is crucial to decipher the ecological aspects of plant biomass degradation. This process ultimately occurs through the central pathways for the metabolism of key aromatic compounds. Countless peripheral pathways converge toward a small number of central intermediates that undergo ring-cleavage in the central pathways. In fungi, there are five main described pathways of the central intermediates: catechol, protocatechuate, hydroxyquinol, homogentisate, and gentisate (2), of which the corresponding catabolic genes are known (3–7). The five central pathways are present in the genomes of both Ascomycota and Basidiomycota, sharing high similarity, despite some noteworthy differences (2). A catabolic pathway for the central intermediate 3-hydroxyanthranilate was recently discovered in fungi (8). The existence of other specific dioxygenases for gallate, hydroquinone, homoprotocatechuate, and pyrogallol, as described for bacteria, remains to be verified in fungi, although mostly accepted that these intermediates are channeled into the five main central pathways (4, 9, 10).

The catabolism of gallate by Ascomycota is assumed to occur through three pathways but none is fully disclosed. First, a path of initial reduction and metabolization to protocatechuate was described in *Penicillium simplicissimum* (9). Second, it was established for the yeast *Blastobotrys adeninivorans* an initial non-oxidative decarboxylation of gallate to pyrogallol that undergoes intradiol ring-cleavage, mediated by a catechol-*like* dioxygenase, forming 2-hydroxymuconate, which is then channeled to an undisclosed catabolic pathway (11, 12). The same pathway was suggested for *Aspergillus* and *Penicillium* species once they also accumulate pyrogallol (13). Finally, direct ring cleavage was proposed for both *A. niger* and *A. flavus*, based on indirect pieces of evidence, including apparent gallate dioxygenase activity (13, 14). The catabolism of gallate in *A. niger* was further described to be independent of the protocatechuate branch of the 3-oxoadipate pathway (15). Collectively, data on the catabolism of gallate in fungi suggest that different pathways may be used by closely related species, and possibly exhibit some redundancy.

To better understand the catabolism of gallate, in the present study, we resorted to *A. terreus* as a model fungus. Specifically, the transcriptional response of the fungus growth on gallate was analyzed by RNA-seq (acetate was used as a control condition). This was complemented by targeted gene expression analyses, functional studies of gene deletion mutants, and analytical chemical identifications of specific intermediates. The data support the elucidation of the catabolism of gallate, and other similar compounds, through a new central pathway with initial hydroxylation (decarboxylating) and formation of the central intermediate 1,2,3,5-tetrahydroxybenzene. Furthermore, genes essential for gallate utilization are organized in a cluster of genes present in several genomes, increasing the relevance of this novel central pathway for fungi in general.

## Materials and Methods

### Strains and growth conditions

*Aspergillus terreus* FGSC A1156 (equivalent NIH2624), *Aspergillus nidulans* FGSC A4, and additional strains (Table S1) asexual spores were harvested and maintained as frozen suspensions at -80 °C (16). Liquid cultures were initiated with 10^6^ spores/mL and incubated with orbital agitation (250 rpm) in the dark at 37 °C. Batch cultivations were performed in 250 mL Erlenmeyer flasks with a working volume of 50 mL. A low nitrogen minimal medium was used containing *per* liter 3 g NaNO_3_, 0.01 g ZnSO_4_·7H_2_O, 0.005 g CuSO_4_·5H_2_O, 0.5 g MgSO_4_·7H_2_O, 0.01 g FeSO_4_·7H_2_O, 0.5 g KCl. Filter sterilized salts were added to an autoclave sterilized 100 mM potassium phosphate solution. The carbon sources were added either directly to the potassium phosphate solution for sterilization (60 mM sodium acetate – control, pH 6.0) or into the mineral media after filter sterilization (*e.g.,* 20 mM gallate, pH 4.0). Whenever relevant, the appropriate auxotroph nutrient requirements were added, and agar (10 g L^-1^) was used to gel the media. Cultures of gene replacement mutants for functional analysis in liquid medium were done in 100 mM potassium phosphate buffer pH 5.0 incubated with orbital agitation (250 rpm) in the dark at 30 °C using pre-grown mycelium for 24 h in sodium acetate liquid minimal medium. **Chemicals.** Gallic acid monohydrate (ACS), 3,4,5-trimethoxyphenol (>98.5%), ascorbic acid (99%), and sodium nitrate (extra pure) were purchased from Acros Organics; 3,4-dihydroxybenzoic acid (97%) from Alfa Aesar; acetonitrile (>99.9%), ethyl acetate (>99.8%), diethyl ether (>99.5%), and methanol (>99.8%) from Fisher Chemical; sodium hydroxide (>98%) from José Manuel Gomes dos Santos; anhydrous sodium sulfate (99%), boron tribromide solution in anhydrous dichloromethane, and anhydrous dichloromethane (99.7%) from Thermo Scientific; agar, and resorcinol (99%) from Panreac; all other chemicals of the highest grade from Sigma-Aldrich/Merck.

### Synthesis of 1,2,3,5-tetrahydroxybenzene

1,2,3,5-Tetrahydroxybenzene was synthesized as previously described [18]. A solution of BBr_3_ in anhydrous CH_2_Cl_2_ (1 M, 18.8 mL) was added, with stirring, to a solution of 3,4,5-trimethoxyphenol in anhydrous CH_2_Cl_2_ (1.88 mmol, 10 mL) at -80 °C under nitrogen during 1 h and the reaction was left to warm until room temperature. After overnight, the mixture was cooled to 0 °C and distilled water (5 mL) was added. The CH_2_Cl_2_ was removed under nitrogen flow and the remaining mixture was extracted with ethyl acetate. The organic phase was dried with anhydrous NaSO_4_, filtered, and concentrated under nitrogen flow. ^1^H NMR (800 MHz, DMSO-*d*_6_): 5.72 (s, 2H); ^13^C NMR (201 MHz, DMSO-*d*_6_): 94.34; 125.33; 146.77; 149.91.

### Metabolites identification and quantification by chromatography

The culture media or the respective ethyl acetate extracts were analyzed by liquid chromatography (LC) for the identification and/or quantification of the aromatic compounds and their transformation products using a previously described method (17). Two other methods were used depending on the polarity of the compounds. For more polar compounds, a Synergi Polar-RP column, 4 μm, 150 x 4.6 mm (Phenomenex), set at 26 °C, was used. For the separation, the eluent (5 mM potassium phosphate pH 3.05 with 2% methanol) was used 6-min under isocratic conditions, followed by a 15 min gradient of increasing percentage of methanol from 2% to 50%, at a flow rate of 0.8 ml·min^-1^ (HPLC method A). For very polar metabolites a Synergi Hydro-RP column, 4 μm, 250 x 4.6 mm (Phenomenex), set at 26 °C, was used. The eluent – 20 mM potassium phosphate pH 2.9, was used under isocratic conditions at a flow rate of 0.7 ml·min^-1^ (HPLC method B).

Samples were analyzed by gas chromatography-mass spectrometry (GC-MS) with an Agilent GC (7820A) equipped with a single quadrupole Agilent (5977B) MS. The ethyl acetate extracts were dried under a soft nitrogen flow, and subsequently derivatized with N,O-bis(trimethylsilyl)-trifluoroacetamide containing 1% of trimethylchlorosilane and pyridine (5:1) for 30 min, 90 °C. The separation of the analytes was carried out using a HP-5MS column (30 m by 250 µm by 0.24 µm; Agilent) with the following ramp temperature: 80 °C, 4 °C/min until 290 °C for 10 min. Full scan mode with a mass range of m/z 40 to 900, with a source at 230 °C and electron impact ionization (EI+, 70 eV), was used for all samples, and acquisition delay was set at 240 s. Technical triplicates were analyzed. Data acquisition was accomplished by MSD ChemStation (Agilent) and compounds were identified based on a spectral library (NIST 2017) and references relying on diagnostic ions distinctive of each derivative and its spectrum profile.

The extracts were also analyzed by liquid chromatography-mass spectrometry (LC-MS) using a Q Exactive Focus Hybrid Quadrupole-Orbitrap (Thermo Scientific). The separation was achieved in a Waters Acquity UPLC HSS T3 column (2.1x150 mm, 1.8 µm particle size), using a gradient of increasing percentage of 0.9% formic acid (FA) in acetonitrile (B) and 0.9% FA in ammonium formate 50 mM (pH 2.9) (C) and decreasing percentage of 0.9% FA in water (A). The flow rate was 0.2 mL·min^-1^ and the column was kept at 30 °C. The sample injection volume was 1 µL. The data were acquired using Xcalibur software v.4.0.27.19 (Thermo Scientific). The method consisted of several cycles of Full MS scans (R=70000; Scan range=75-1125 m/z) followed by 3 ddMS2 scans (R=17500; 20, 40, 60 NCE) in negative mode. External calibration was performed using LTQ ESI Negative Ion Calibration Solution (Thermo Scientific). The generated mass spectra were processed using Compound Discoverer 3.2 (Thermo Scientific) for small molecule identification.

### Nuclear magnetic resonance spectroscopy

Ethyl acetate extracts were dried under a soft nitrogen flow (approx. 5 mg) and solubilized in 500 μL of DMSO-*d*_6_ for NMR analyses. All NMR spectra (^1^H, ^1^H−^1^H COSY, ^1^H−^13^C HSQC, ^1^H−^13^C HMBC) were acquired at 25 °C using an Avance III 800 CRYO spectrometer (Bruker Biospin, Rheinstetten, Germany) in 5 mm diameter NMR tubes. MestReNova, Version 11.04-18998 (Mestrelab Research, S.L.) was used to process the raw data.

### Transcriptome Analysis

Total RNA extraction of liquid-grown mycelia was performed essentially as previously described (16), using a Tissuelyser LT (Qiagen) for cell disruption. The quality and quantity of RNA were determined by capillary electrophoresis using an HS RNA kit and 5200 Fragment Analyzer (Agilent). For single-end RNA sequencing (RNA-seq), libraries were generated using the Smart-Seq2® mRNA assay (Illumina, Inc.) according to the manufacturer’s instructions. Samples were indexed and sequenced on the Illumina NextSeq550 (20M reads *per* sample). Generated FastQ files were analyzed with FastQC and any low-quality reads were trimmed with Trimmomatic (18). All libraries were aligned to *A. terreus* NIH2624 genome assembly (ASM14961.v1) with gene annotations (built 2010-06) from Ensembl Fungi using HISAT2 v. 2.2.1 (19) with sensitive option and maximum intron length at 4000. Read counts were obtained using featureCounts v2.0.1 (20). All RNA-seq experiments were carried out in three biological replicates. Differential expression analysis was performed using DESeq2 v.1.24.0 (21). Transcript abundance was defined as transcripts *per* million (TPM). The genes that showed more than log_2_ 1-fold expression changes and above median TPM with *p*-adj value <0.05 are defined as significantly differentially expressed genes in this analysis.

Two strategies were implemented to identify upregulated genes that were absent or incorrectly annotated in the current *A. terreus* NIH2624 genome assembly. To identify genes that could be incorrectly joined together in the annotation, the differentially expressed exons were determined as described above for gene features. Genes that showed simultaneously up- and downregulated exons were manually curated. To identify genes that were not in the current annotation, the entire genome was divided into 500 bp length sections, which were analyzed as described above as if they were genes/exons. The upregulated differentially expressed sections (threshold set for upper quartile expression levels) not attributed to a current annotated gene were manually curated. Automated gene structure prediction of new genes or poorly annotated genes was done using AUGUSTUS (22). Gene structure was manually curated according to RNA-seq data and used in the transcriptome analysis (Table S2).

Protein domain entries were obtained from InterPro 90.0 either from predetermined data or using InterProScan (REST) service to search entries for the new gene predictions (23). The enrichment analyses were performed with the FunRich software v.3.1.3 (24), with a *p*-value <0.05 after Bonferroni correction. Functional characterization and metabolic pathways analysis was performed using the KEGG application KofamKoala (25). Metabolic gene clusters were identified within annotated genomes through sequence homology searches using cblaster v.1.3.12 (26).

### Reverse transcription-quantitative PCR analysis (RT-*q*PCR)

Total RNA was extracted as reported above, and cDNA synthesis followed a previously described protocol (16). Oligonucleotide pairs were designed using Primer-BLAST (27) and supplied by STAB Vida (Oeiras, Portugal) (Table S3). The RT-*q*PCR analysis was performed in a CFX96 Thermal Cycler (Bio-Rad), using the SsoFast EvaGreen Supermix (Bio-Rad), 250 nM of each oligonucleotide and the cDNA template equivalent to 10 ng of total RNA, at a final volume of 10 µl per well, in three biological replicates. The PCR conditions were: enzyme activation at 95 °C for 30 s; 40 cycles of denaturation at 95 °C for 5 s, and annealing/extension at 60 °C for 15 s; and a melting curve obtained from 65 °C to 95 °C, consisting of 0.5 °C increments for 5 s. Data analyses were performed using the CFX Manager software v.3.1 (Bio-Rad). The expression of each gene was taken as the relative expression in pair-wise comparisons of each condition relative to the acetate control. The expression of all target genes was normalized by the expression of the gamma-actin gene ATEG_06973 or AN6542.

### Generation of gene replacement mutants

The genes AN0331, *hmg*A (AN1897), AN4576, AN10530, and AN12484 were selected for gene replacement with *A. fumigatus pyr*G in *A. nidulans* A1149 strain, performed according to a well-established method (28), essentially as previously described (3). Oligonucleotides were designed as described above (Table S3). Gene replacement was confirmed by PCR (Fig. S1).

### Data availability

All relevant data is available in the manuscript. Supplementary figures and tables are provided to support the results presented on this manuscript. Gene transcript TPM values are contained in Table S4, and raw data with metadata were deposited in the Sequence Read Archive under Bioproject accession number PRJNA612036.

## Results and discussion

### During gallate catabolism in *A. terreus,* the 3-oxoadipate and homogentisate central pathways are mostly unregulated

A comparative transcriptome analysis was performed here aiming to disclose the central pathways relevant to the catabolism of gallate in *A. terreus*. During gallate catabolism (approx. 30% consumed of 20 mM), 1545 encoding genes were found differentially upregulated and 1756 downregulated (see Fig. S2 for principal component analysis and MA plots, and Table S4). Functional analysis of differentially expressed genes revealed that the global response to gallate in comparison to the control acetate differed in the regulation mechanisms (Fig. S3). Gallate-induced response mainly occurs at the transcriptional regulation, and resorts more to fungi-specific DNA binding transcription factors (IPR036864). On the contrary, acetate-induced response apparently occurs more at the translational and post-translational regulation and involves proteins containing RNA binding/modifying domains (IPR012340, IPR012677, IPR001412, IPR014729, IPR009000) and proteasome degradation domains (IPR001353, IPR029055, IPR023332). This may however be biased since more is unknown in the upregulated genes dataset. About 21% of upregulated genes don’t have a known protein domain in comparison to only *ca.* 10% in the control (17.6% in all genes). Moreover, 72% of upregulated genes are currently not functionally annotated using KEGG, while only 36% of the downregulated ones are not. The Major Facilitator Superfamily (MFS) of transporters (IPR036259) was 2-fold enriched and represented 8% of all upregulated genes. The general response to gallate is relatively similar to that towards salicylate (*e.g.* enriched protein domain families) (8), except for the secondary metabolism which is mostly unregulated in gallate but upregulated in salicylate.

The known central pathways, namely the 3-oxoadipate, the homogentisate, and the 3-hydroxyanthranilate pathways for the catabolism of aromatic hydrocarbons were either unregulated or showed upregulation with moderate transcripts abundance (Table S5). The 3-oxoadipate pathway was globally the central pathway most responsive to gallate, especially the protocatechuate branch (Table S5). Quinate or shikimate catabolism (or hypothetically from an initial reduction of gallate) via protocatechuate is dependent on *qut*C (ATEG_00348), 3-dehydroshikimate dehydratase, which mediates the reaction that connects the upper and the central catabolic pathways (*e.g.* protocatechuate branch), intersecting also with the anabolic shikimate pathway (29). The predicted quinate utilization cluster (ATEG_00348 to ATEG_00355), including *qut*C, was scarcely expressed. Similarly, a second quinate utilization-*like* cluster (ATEG_07672_new to ATEG_07676), not present in *e.g.*, *A. nidulans*, was also scarcely expressed (Table S5). Other *qut*C homologs were unregulated, nearly excluding a path of initial reduction of gallate. Moreover, as previously reported for *A. niger* (15), we also observed that the protocatechuate branch is not essential for the catabolism of gallate in *A. nidulans* (see below).

### Four genes compose the gallate catabolic pathway in *A. terreus*

The transcriptional reprograming induced by gallate was confirmed by the upregulation and very high expression of several tannase genes: ATEG_02651 orthologue of AO090023000047 (30), ATEG_03047 orthologue of An18g03570 (31), and ATEG_07937 orthologue of AO090103000074 (32). The tannic acid transcriptional activator-repressor module composed of *tan*R (ATEG_10383) and *tan*X (ATEG_10381) (33) was also found to be upregulated though more modestly expressed. Additional esterase genes (ATEG_08115b_new and ATEG_09944) and *O*-methyltransferases (ATEG_07862, ATEG_09300, and ATEG_01928) were also highly expressed and upregulated, suggestive of a role in the metabolism of esters of gallate or their precursors (Table S5).

Some gene clusters of upregulated genes were identified but none contained the abovementioned tannases, esterases, and *O*-methyltransferases. The central pathways for the catabolism of aromatic compounds are every so often organized into clusters of genes in the genomes of fungi (2). The hypothesis that some gallate upregulated genes in *A. terreus* are in a gene cluster in other species was tested using the top 50 abundant transcripts with an FC>2 (Fig. 1; similar results were obtained using the top 100). Three main groups of gene clusters were identified; these comprise (i) sugar/inositol transporters genes (IPR003663), (ii) genes of the ethanol utilization pathway (discussed below), and (iii) four genes, including a putative monooxygenase and a fumarylacetoacetate hydrolase (FAH)-*like* gene – hallmarks of a catabolic gene cluster of aromatic compounds. Further analysis of the last main group (iii) revealed that three gene cluster types, each having a distinct monooxygenase as a core gene, are present across species of Ascomycota (Fig. 2) and only one in Basidiomycota (Fig. S4). The genes of the three cluster types are in four loci in the genome of *A. terreus,* and four of them highly expressed and upregulated (ATEG_04855, ATEG_04856, ATEG_05726b_new, and ATEG_08221-22_new; hereafter denominated the four catabolic genes of the gallate pathway) (Fig. 3 and Table S5). Two of the three monooxygenase genes were found upregulated, though ATEG_05726b_new was more highly expressed. The encoded protein of the *A. niger* ortholog (NRRL3_4659), can convert *in vitro* protocatechuate in 1,2,4–trihydroxybenzene (hydroxyquinol), hence recently suggested to convert gallate to 1,2,3,5–tetrahydroxybenzene (34). The other highly expressed genes present in the three types of clusters are a FAH-*like* (ATEG_04856), a fungal conserved short protein gene (ATEG_04855) with no known domains (containing a conserved nucleophilic residue serine, a.a. 64, and only found in Dykaria), and a DUF3500 gene (ATEG_08221-22_new). In bacteria, DUF3500 genes are also often located adjacent or in the vicinity of a FAH-*like* gene not closely related to the characterized ones (data not shown). They are mostly only present in FCB, Proteobacteria, PVC and Terrabacteria groups, and Acidobacteriota phylum. Finally, in fungi, DUF3500 genes are mostly found in filamentous Ascomycota (*i.e.* Pezizomycotina subphylum) and Basidiomycota with macroscopic fruiting bodies (*i.e.* Agaricomycotina subphylum) (23).

**Figure 1.**
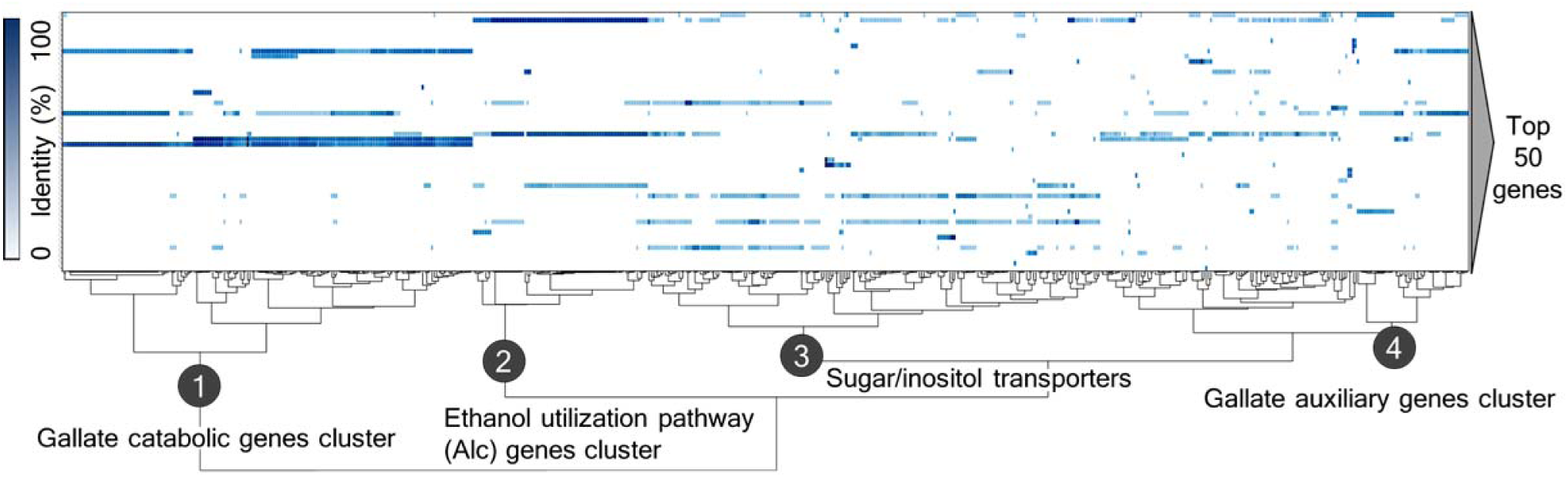
Gene cluster analysis with the top 50 most abundant genes (TPM) differentially upregulated (Fold Change>2) in response to gallate. Co-localized genes were identified within GenBank annotated genomes through sequence homology searches, and their hierarchical clustering analysis done using the program cblaster (603 clusters across 578 genomic scaffolds from 375 organisms were detected; see also Table S6). Four main clusters were observed including one containing the four catabolic genes of the gallate pathway.

**Figure 2.**
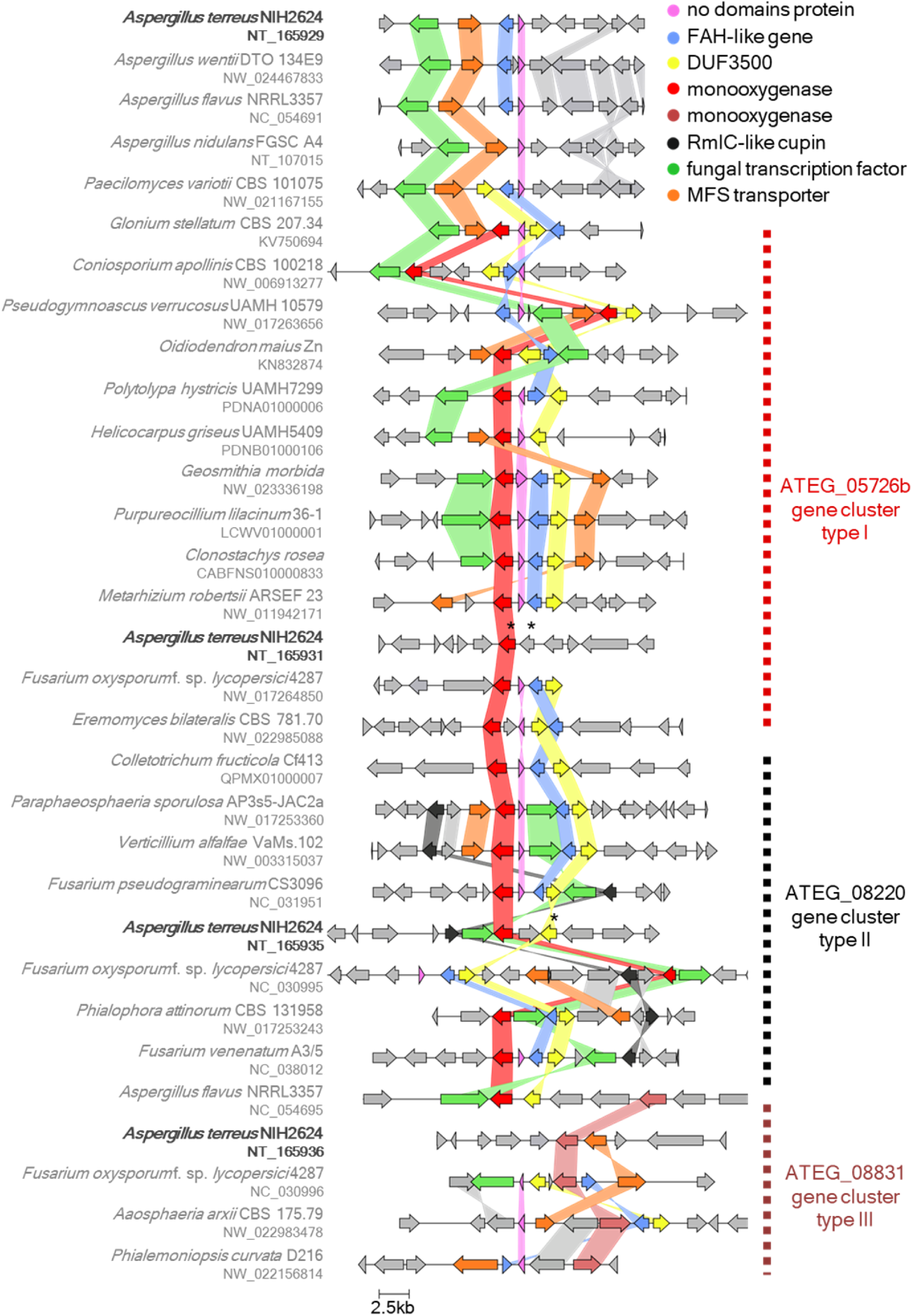
Gallate and related gene clusters identified in Ascomycota fungi. Here are depicted selected examples of the three main gene cluster types having a distinct monooxygenase as a core gene (highlighted by the *Aspergillus terreus* genes ATEG_05726b, ATEG_08220, and ATEG_08831). These gene clusters additionally often include a fumarylacetoacetate hydrolase (FAH) *like* gene (ATEG_04856), a DUF3500 gene (ATEG_08221-22), and a fungal conserved short protein with no known domains’ gene (ATEG_04855) alongside with transporter (MFS) and transcription factor genes. Clusters were identified within GenBank annotated genomes through sequence homology searches and graphical representation using the programs cblaster and clinker, respectively. Homologous genes are represented using a color code and gene links represent an identity threshold of over 30%. The locus accession number is given below the species name and the asterisk indicates a gene with a new structure annotation.

**Figure 3.**
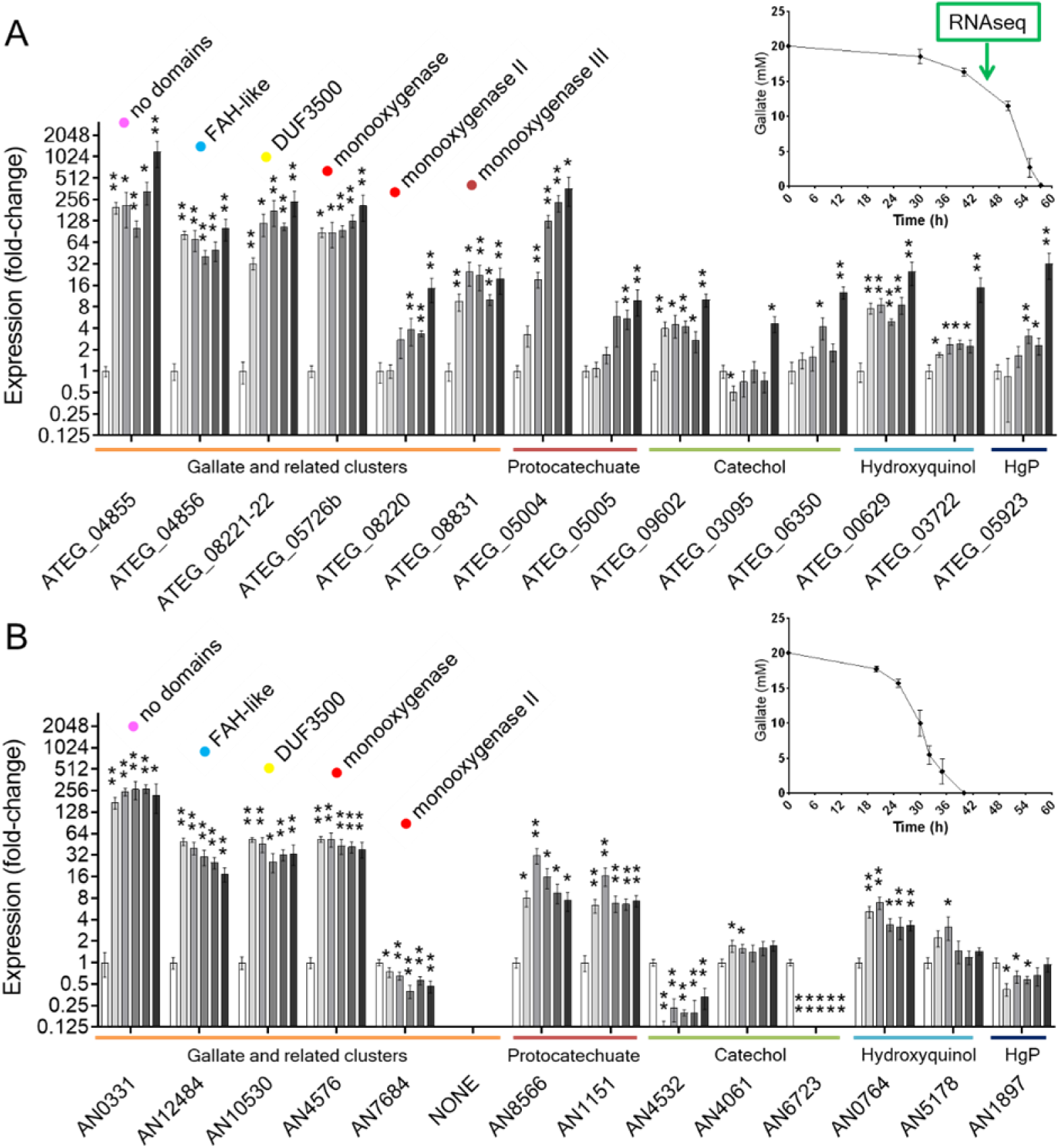
Gene expression of the gallate pathway genes and other central pathways of *Aspergillus terreus* (panel A) and *A. nidulans* (panel B) along the time of cultivation in gallate media (grey bars) compared to the control in acetate media (white bar). The distinct monooxygenase types from gallate-*like* gene clusters are also shown. Representative genes of the 3-oxoadipate pathway (protocatechuate and catechol branches and the hydroxyquinol variant) and the homogentisate pathway (HgP) were chosen including the respective dioxygenase gene. The timepoints for gene expression in gallate media correspond to those depicted in the graphical inserts for the consumption of gallate (* *p*-value < 0.05; ** *p*-value < 0.005).

The massive expression and upregulation of the ethanol utilization pathway (Table S5) constituted of alcohol dehydrogenase *alc*A (ATEG_09407 orthologue of *alc*A/AN8979), aldehyde dehydrogenase *ald*A (ATEG_05020 orthologue of *ald*A/AN0554) and transporter *alc*S1 (ATEG_03626-27_new; with homology to *alc*S/AN8981 and AN8390 (35)) strongly indicates the production of their physiological inducer acetaldehyde (35). The regulator ATEG_03627 with homology to *alc*R/AN8978 was also positively regulated. Acetaldehyde can be produced from the catabolism of compounds as diverse as ethanol, amino acid threonine, catechol (meta-cleavage), or phenylpropionic acids (aka hydroxycinnamic acids). Cytosolic pyruvate carboxylase *pyc*A (ATEG_05433) was found to be upregulated, most likely contributing to the formation of acetaldehyde and induction of the ethanol utilization pathway. Additionally, *acu*A (acetyl-CoA synthetase), *acu*C (transcriptional activator), and *cp*A (transport of acetate ions with homology to *alc*S) were upregulated despite the control condition acetate.

The four catabolic genes of the gallate pathway, but not the ethanol utilization pathway, were also found upregulated in *A. flavus* in the presence of gallate, notwithstanding the co-metabolic conditions used in this previous study (36).

### The four catabolic genes of the gallate pathway are essential for gallate utilization in *Aspergillus*

The transcriptional gallate-induced response suggests that the main catabolic pathway for gallate in *A. terreus* starts with initial hydroxylation (decarboxylating) and formation of the central intermediate 1,2,3,5–tetrahydroxybenzene. Processing of this intermediate through the central pathway results in the buildup of end metabolites that trigger the induction of the ethanol utilization pathway and C2 metabolism. To confirm this hypothesis, gene deletion mutants were produced in *A. nidulans,* and gene functional assays were performed. Gene expression analysis first validated the results obtained in RNA-seq of *A. terreus* (Fig. 3A). A similar gene regulation was observed for *A. nidulans* mostly verifying that the same pathway is used in both species (Fig. 3B). The four catabolic genes of the gallate pathway showed a remarkable induction from the early culture timepoints, in opposition to the other genes of the central pathways analyzed.

Moreover, the four catabolic gene deletion mutants showed severely impaired growth in gallate and tannic acid solid media (Fig. 4) demonstrating their essential role in the catabolism of gallate. Growth of deletion mutants in other aromatic compounds, namely 2,3-dihydroxybenzoate, phenylacetate, phloroglucinol, protocatechuate, and resorcinol was unaffected (Fig. 4 and S5). This revealed that the four catabolic genes are most likely only essential to the catabolism of gallate and other compounds that share the same central pathway unrelated to those previously described. Growth in an isomer of gallate, 2,4,6-trihydroxybenzoate, showed severe growth impairments like those imposed by gallate except for the monooxygenase deletion-mutant (ΔAN4576). This indicated that the central metabolite 1,2,3,5-tetrahydroxybenzene was formed through hydroxylation (decarboxylating) of gallate and its isomer.

**Figure 4.**
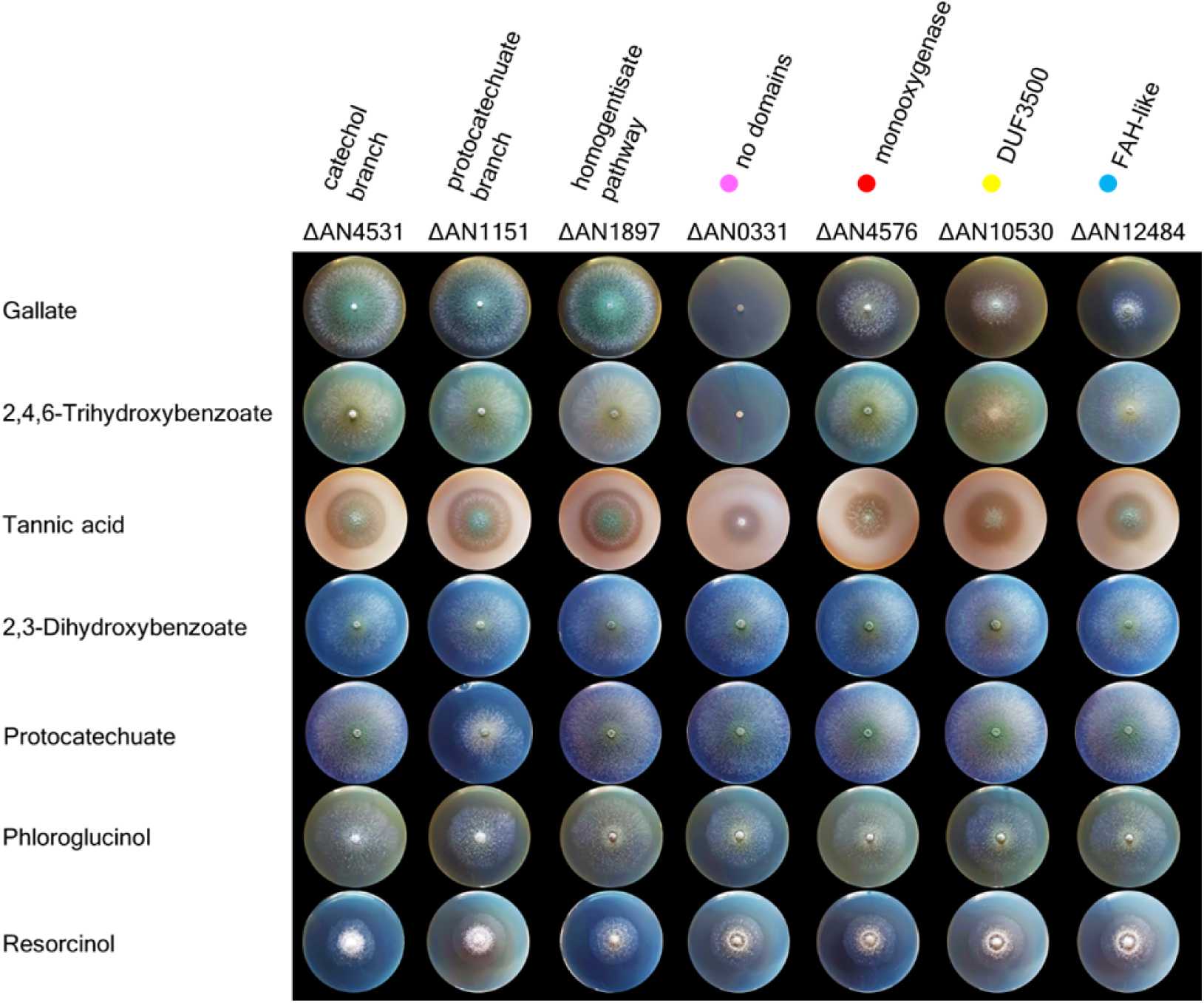
*Aspergillus nidulans* gene deletion mutants grown in gallate and other carbon sources. Mutant strains of the four gallate pathway genes showed phenotypes of growth impairment for gallate, its isomer 2,4,6-trihydroxybenzoate (except for ΔAN4576), and tannic acid utilization as a carbon source. ΔAN10530 growth media also accumulated a colored metabolite for gallate or related compounds. The mutants used as control namely of the catechol branch or protocatechuate branch of the 3-oxoadipate pathway, and homogentisate pathway showed the expected growth phenotypes in the respective metabolites of their pathways, *i.e.*, normal growth for ΔAN4531 in 2,3-dihydroxybenzoate, growth impairment for ΔAN1151 in protocatechuate (note: phenylacetate was a very poor carbon source for all strains at the low pH used). Aromatic hydrocarbons (20 mM) or tannic acid (2.5 mM) were added as the main carbon source to solid minimal media (pH 5.0).

### 1,2,3,5–Tetrahydroxybenzene is the central intermediate of the gallate catabolic pathway in *Aspergillus*

To further validate 1,2,3,5–tetrahydroxybenzene as a new central intermediate, the metabolites accumulating after the addition of gallate or its isomer to acetate pre-grown cultures of gene deletion mutants were analyzed. Nor gallate or their metabolites accumulated in incubations of the Δ gallate 1-monooxygenase (ΔAN4576), suggestive of functional redundancy (Fig. S6 and S7). Transient accumulation of 1,2,3,5–tetrahydroxybenzene in cultures of gallate and its isomer was observed for the Δ DUF3500 gene (ΔAN10530) in the presence of supplemented ascorbic acid (Fig. 5 and S8). 1,2,3,5–Tetrahydroxybenzene was shown to be unstable in phosphate buffer even at lower pH, most likely due to autooxidation. The resulting autooxidation metabolites were mostly not identified by GC-MS except for 2–furoic acid (Fig. 5 and S10). Accumulation of a single compound was detected in Δ no domains short protein (ΔAN0331); identified as 5-hydroxydienelactone based on NMR and mass spectral data (Fig. 5, 6, and S9). For the Δ FAH-*like* gene (ΔAN12484), the transient accumulation of a major unidentified compound (calculated MW 223.04795 and putative formula C_10_H_9_NO_5_ by LC-MS; non-aromatic by NMR; resolved as three analytes with the same molecular ion by GC after derivatization) and some other metabolites including acetopyruvate was noticed (Fig. 5, S11, S12 ant Table S7). FAH is a diversified superfamily of proteins (IPR011234) with catalytic functions of hydrolase, lyase, and isomerase (37). The nature of the molecule that transiently accumulated in cultures of this gene deletion mutant remains unresolved. It is also not possible to state whether it is a direct or indirect product, the latter being the result of redundant enzymatic activities. However, it may be hydrolyzed to form, at least non-specifically, acetopyruvate or a reduced form (putative 2-hydroxy-4-oxopentanoic acid) as these were only identified in Δ FAH-*like* cultivations (Fig S11).

**Figure 5.**
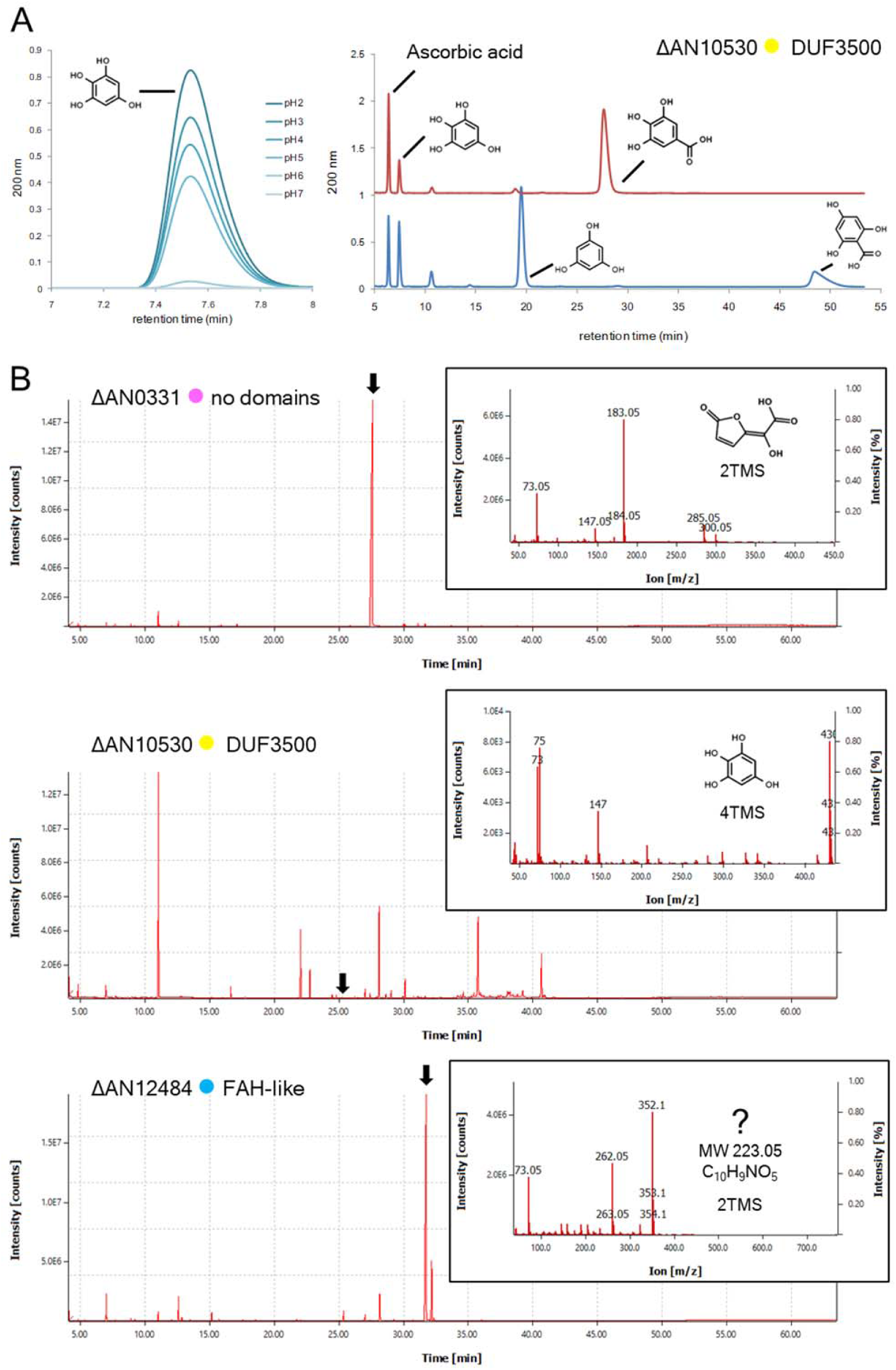
Gallate-derived metabolites differentially accumulated in cultures of the four catabolic gene deletion mutants. HPLC chromatograms (**panel A**) of 1,2,3,5-tetrahydroxybenzene in different pH (10 mM potassium phosphate buffer) for 30 min (left side) and its accumulation in ΔAN10530-DUF3500 cultures (16h, pH 3.0) in the presence of ascorbic acid from either gallate or its isomer 2,4,6-trihydroxybenzoate (right side; HPLC method B). GC-MS chromatograms (**panel B**) for the respective gene replacement mutants’ culture extracts and the mass spectrum of selected analyte molecules (indicated by an arrow).

**Figure 6.**
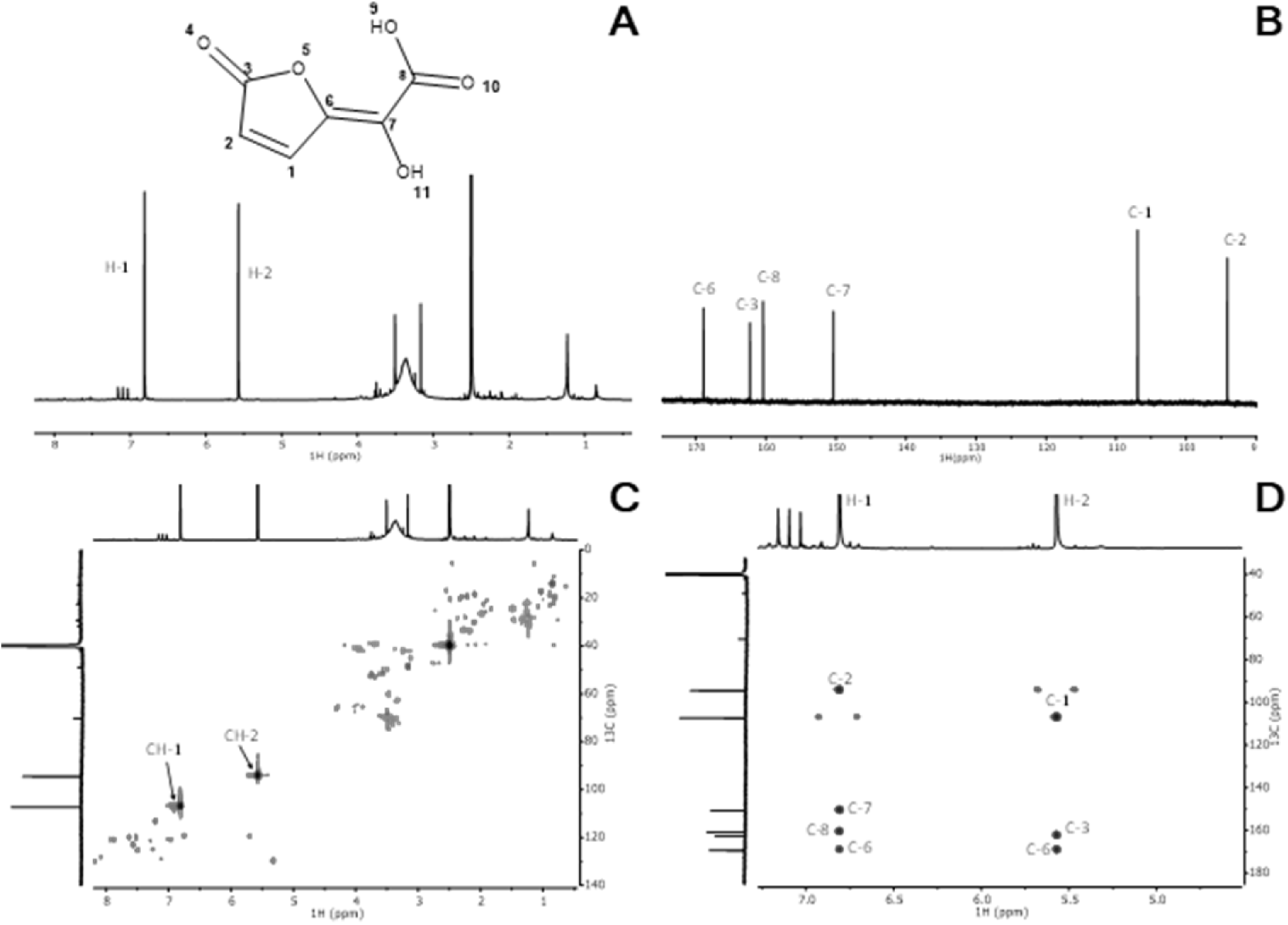
Wide-ranging NMR spectral characterization of ΔAN0331 sample extract with the assignment of the signals related to the presence of 5-hydroxydienelactone. The ^1^H NMR (A); The ^13^C NMR (B); the ^1^H-^13^C HSQC spectrum (C) and the ^1^H-^13^C HMBC spectrum (D). **5-hydroxydienelactone:** ^1^H NMR (800 MHz, DMSO-*d*_6_): 5.57 (d, J=2.2 Hz, 1H); 6.81 (d, J=2.2 Hz, 1H); ^13^C NMR (201 MHz, DMSO-*d*_6_): 94.11; 106.92; 150.37; 160.38; 162.28; 168.93.

In summary, the proposed pathway starts with an initial hydroxylation (decarboxylating) step, which is followed by ring-cleavage (di)oxygenation and immediate (spontaneous) cyclization to create 5-hydroxydienelactone. This compound is then activated to form a resulting nitrogenous adduct and ultimately hydrolyzed to produce acetopyruvate or its reduced form (Fig. 7).

**Figure 7.**
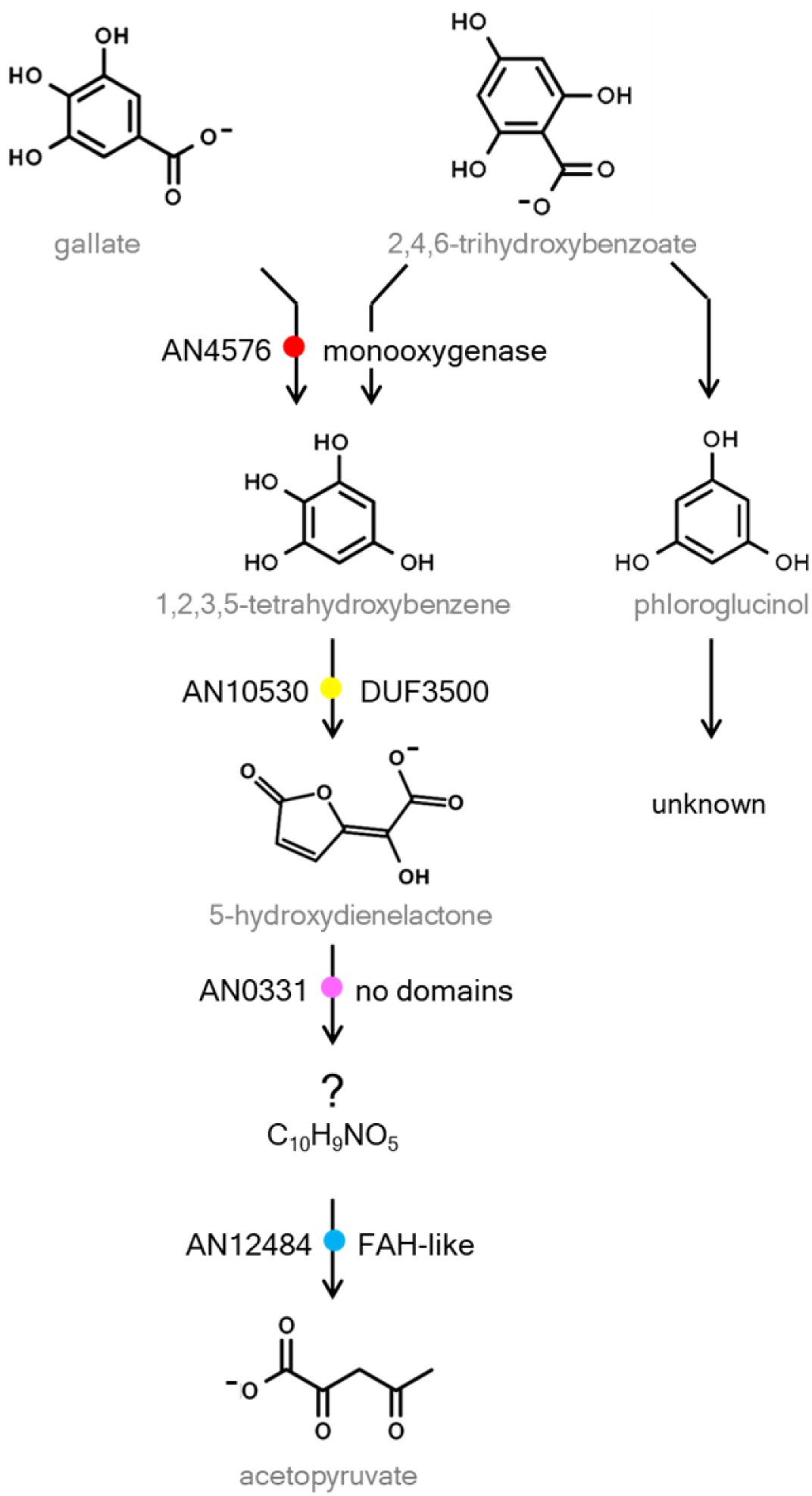
The proposed gallate and related aromatic compound catabolic pathway through the central intermediate 1,2,3,5-tetrahydroxybenzene.

## Conclusions

In this study, we report the discovery of a novel central pathway in fungi responsible for the catabolism of gallate and related aromatic compounds, using the central intermediate 1,2,3,5-tetrahydroxybenzene (Fig. 7). Surprisingly, this pathway differs from those described and/or proposed before for the catabolism of gallate in both bacteria and fungi (38). Based on this discovery, it is wrong to refer to gallate as a central metabolite for fungi; instead, gallate is one of several compounds channeled toward this novel central pathway. The pathway contains two genes encoding proteins having yet no known activities (*i.e.*, DUF3500 and no domains short protein), which necessitates further investigation into their unusual mechanisms of action. However, based on the data, we hypothesize that the DUF3500 protein functions as a newly discovered ring-cleavage enzyme.

The intermediate 1,2,3,5–tetrahydroxybenzene was previously reported in the catabolism of aromatic compounds in bacteria, specifically in the catabolism of gallate or phloroglucinol. It functions as a cocatalyst in the anaerobic conversion of pyrogallol to phloroglucinol, *i.e*., after initial non-oxidative decarboxylation of gallate (39), or as a hydroxylation intermediate in the catabolism of phloroglucinol (40). Only the last partially resembles the central pathway reported here, as acetopyruvate was also identified as a near-end-metabolite. They however may differ, because in the bacteria phloroglucinol catabolism proceeds through this pathway but this dependence was not observed here in *A. nidulans* (Fig. 4). In a recent study, the derivative 6-methoxy–1,2,4–trihydroxybenzene (6-MeOTHB) of the central intermediate 1,2,3,5– tetrahydroxybenzene was identified during the catabolism of syringate (also, a methoxy derivative of gallate) in the Basidiomycota *Phanerochaete chrysosporium* (41). 6-MeOTHB was converted *in vitro*, by hydroxyquinol 1,2–dioxygenase, to 4-hydroxy–2–methoxy–*cis*,*cis*– muconate (41) but the subsequent catabolic steps were unlooked. The DUF3500 and the no domains short protein genes are both absent in *Phanerochaete* genomes, despite being present in genomes of various species in the subphylum Agaricomycotina. These findings suggest that additional pathways for the metabolism of 1,2,3,5–tetrahydroxybenzene or its methoxy derivatives await discovery. Finally, the DUF3500 gene is well represented in certain bacteria genomes, including for example *Sphingomonas*, where it may explain the redundancy observed in this bacteria; deletion of the known key gallate catabolic genes - LigAB (protocatechuate 4,5-dioxygenase), DesZ (3-*O*-methylgallate 3,4-dioxygenase) and DesB (gallate dioxygenase) - did not block gallate utilization (42).

The newly discovered 1,2,3,5–tetrahydroxybenzene central pathway is common in certain subphyla of fungi and likely plays a significant role in the breakdown of aromatics, particularly those derived from syringyl-rich lignin, including gallate and related compounds. Our findings significantly expand the current understanding of the catabolism of aromatic compounds in fungi. This information is crucial for comprehensive studies on the degradation of complex polymers like lignin, and surely may impact the development of technologies aimed at deconstructing and converting them into added-value chemicals.

## Acknowledgements

Mass spectrometry data were generated by the Mass Spectrometry Unit (UniMS), ITQB/iBET, Oeiras, Portugal. All members of the Silva Pereira lab are also thanked for useful discussions.

## Ethics Approval Statement

not applicable

## Supplemental Figures

**Figure S1.**
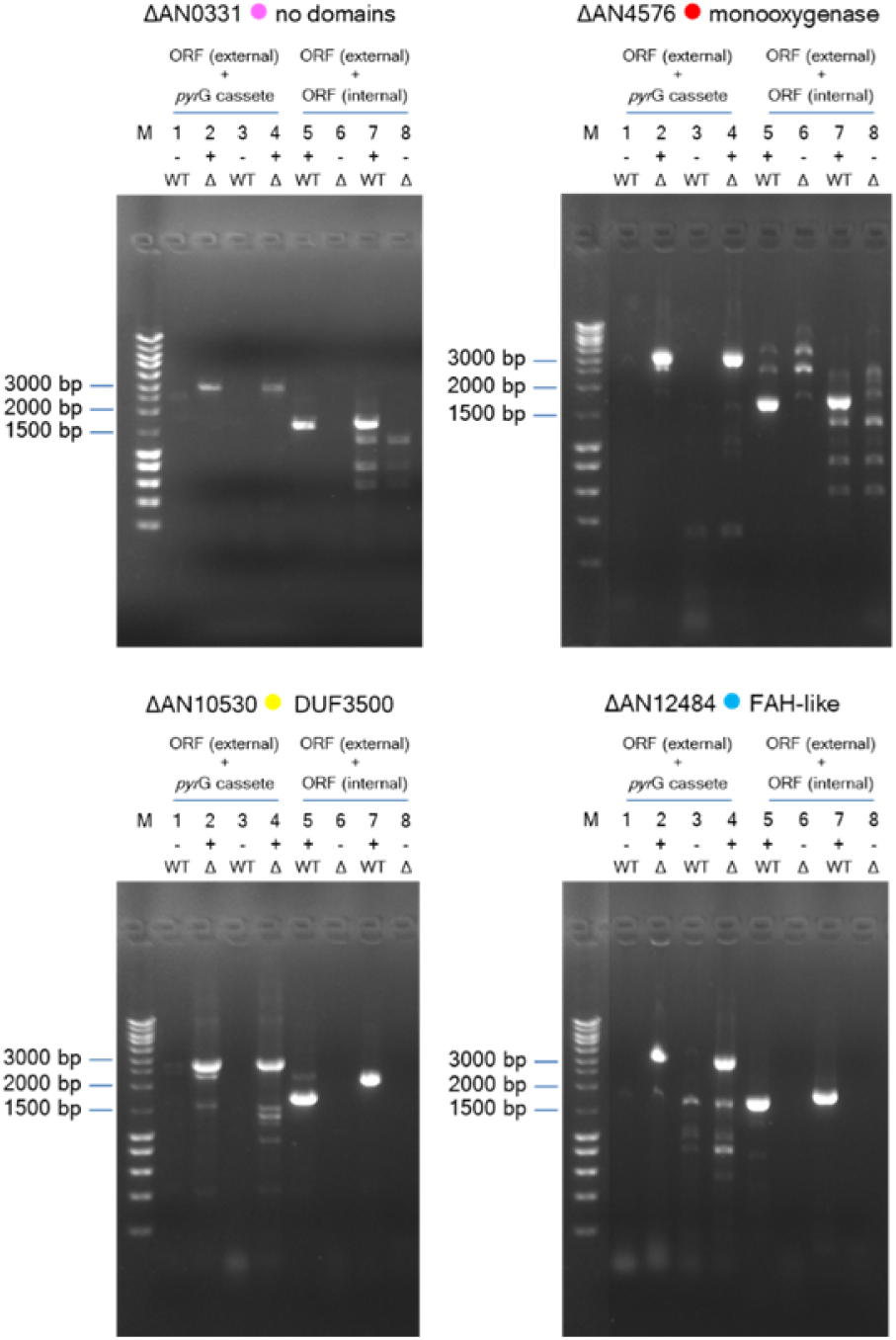
Confirmation of gene replacement mutants by PCR. Mutants (Δ) in opposition to wild type (WT) are positive for amplification using oligonucleotides for both ORF (external) and transformation cassette containing *pyr*G^Afu^ (±2800 bp) and are negative for amplification using oligonucleotides for both ORF (external) and ORF (internal) (1500 to 2000 bp). M - NZYDNA Ladder III. 1 and 2 – oligonucleotide gene_replacement_P1 and *pyr*G^Afu^ cassette amplification reverse. 3 and 4 – oligonucleotide gene_replacement_P6 and *pyr*G^Afu^ cassette amplification forward. 5 and 6 – oligonucleotide gene_replacement_P1 and RT-*q*PCR reverse. 7 and 8 – oligonucleotide gene_replacement_P6 and RT-*q*PCR forward. Even lanes with gene deletion mutant (Δ) genomic DNA and odd lanes with wild-type genomic DNA.

**Figure S2.**
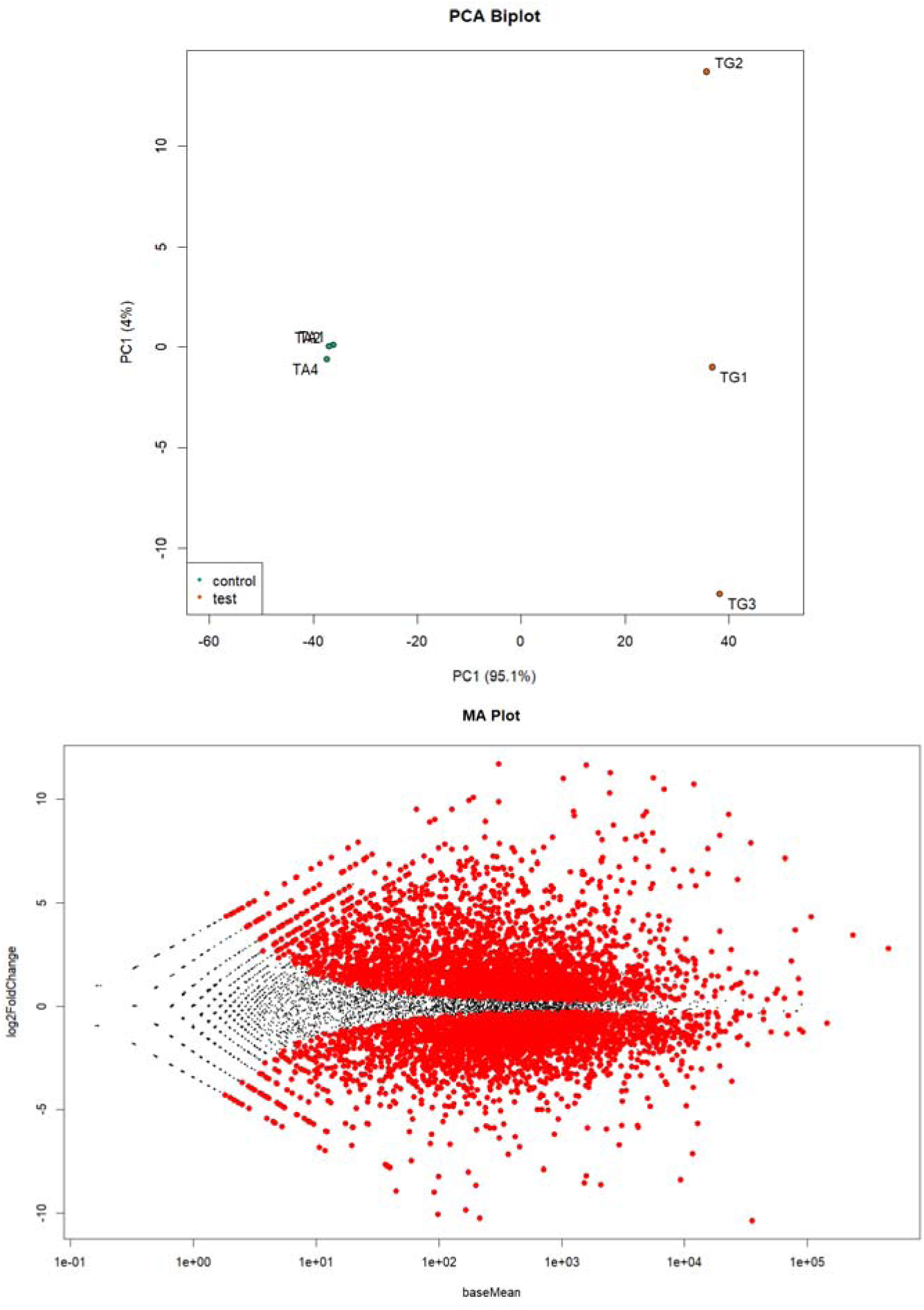
Baseline statistical representation of the RNA-seq data collected for *Aspergillus terreus* samples. The Principal Component Analysis (top) shows the distribution of the whole-transcriptome reads of each of the analyzed replicas (A - acetate medium; G - gallate medium). In the MA plot (bottom) is possible to visualize the differences between the measurements taken between the expression of genes in acetate medium and gallate medium. Both plots were computed using the tools embedded in the DEseq2 package.

**Figure S3.**
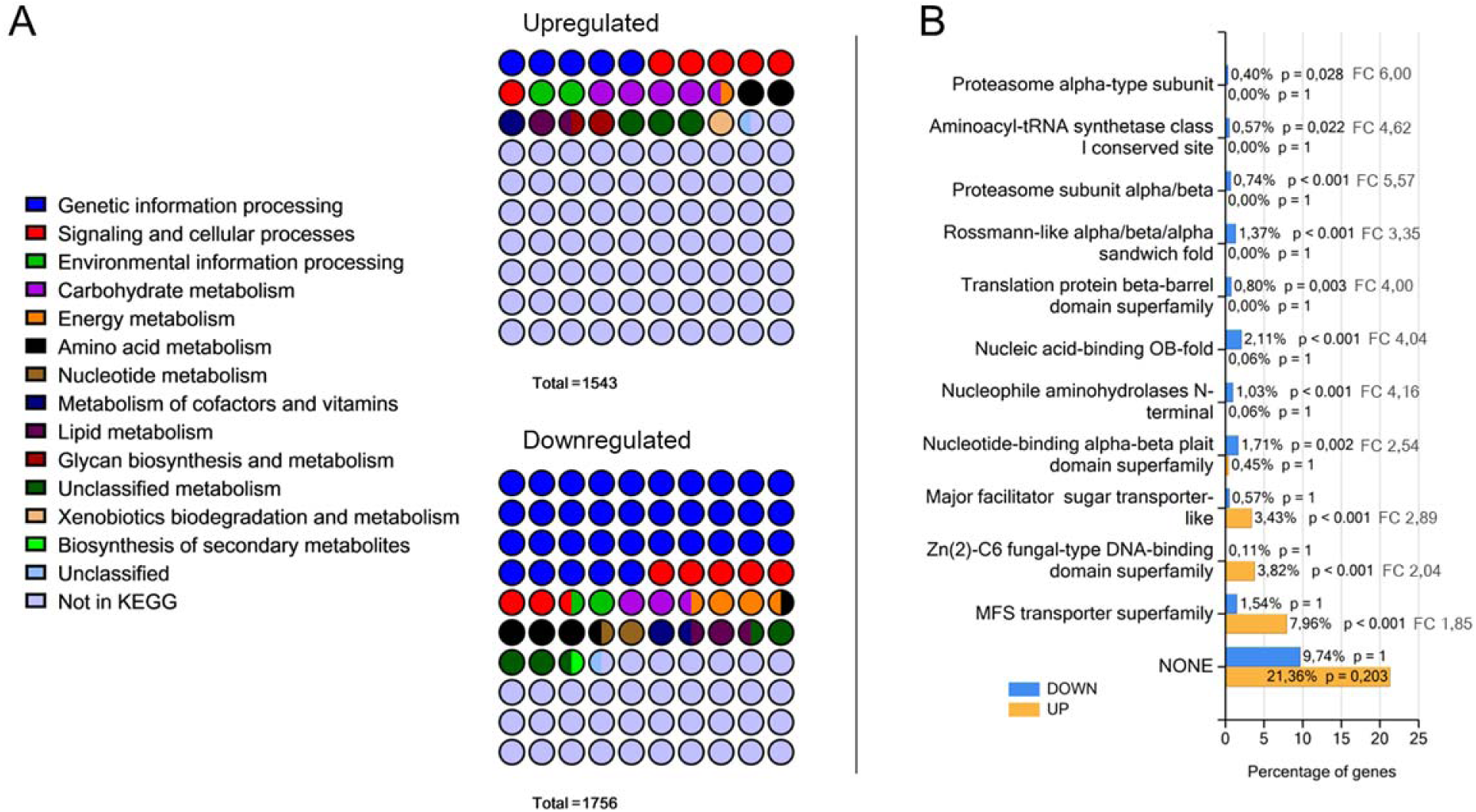
Analysis of the RNA-seq data collected for *Aspergillus terreus* grown in gallate (compared to acetate) comprising functional characterization and protein domains enrichment of the observed differentially expressed genes. KEGG functional categories were slightly simplified by creating super categories that merge the most similar ones. The predicted non-redundant protein domains of the significantly enriched genes in gallate (*p*-value ≤ 0.05) are shown. Percentage of genes are displayed on the bottom axis and the fold change (FC) and log_10_ (*p*-value) next to each bar.

**Figure S4.**
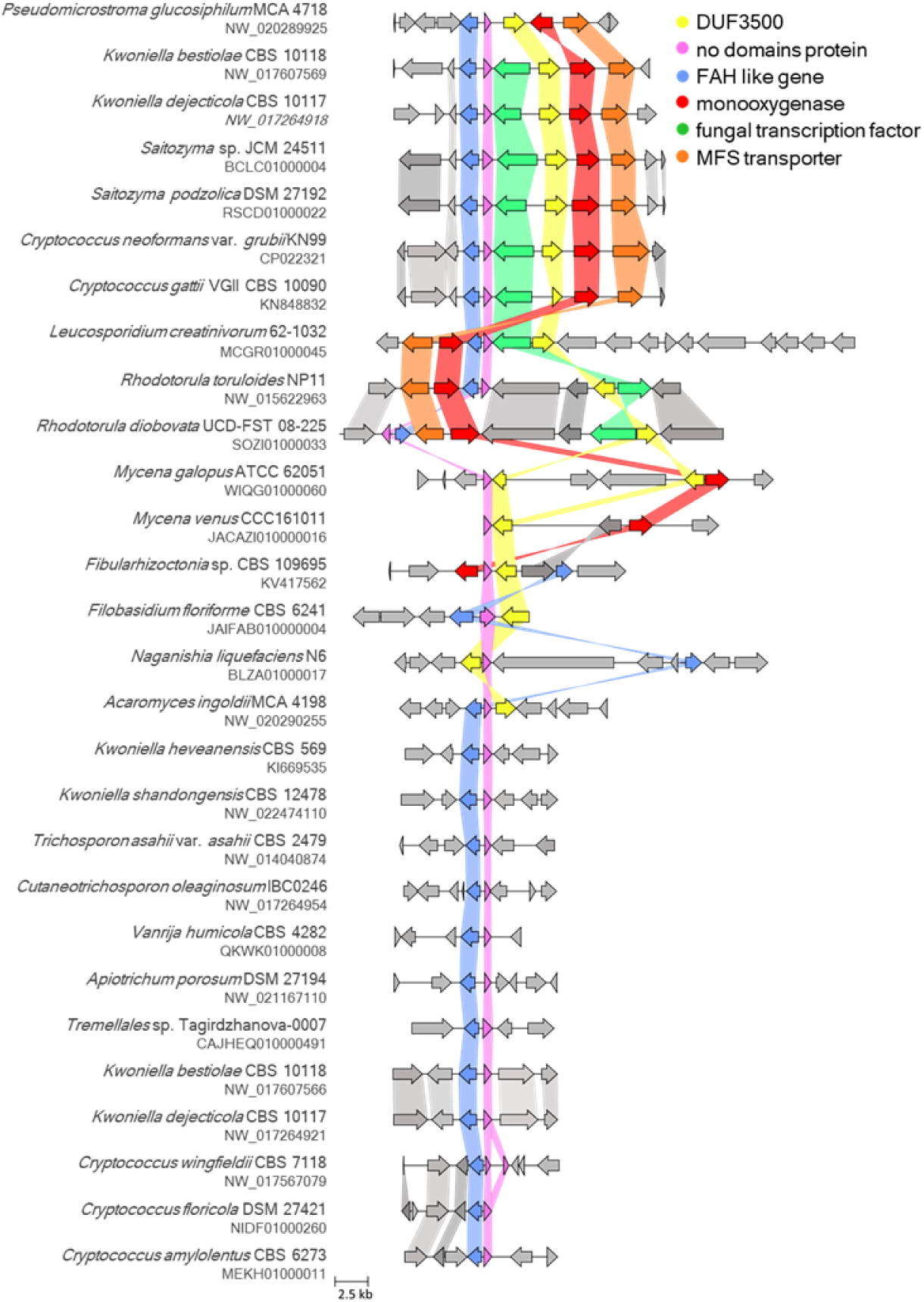
Gallate and related gene clusters identified in Basidiomycota fungi. Here are depicted selected examples of the Basidiomycota gene cluster type often having a monooxygenase, a fumarylacetoacetate hydrolase (FAH) *like* gene, a DUF3500 gene, and a fungal conserved short protein gene alongside transporter (MFS) and transcription factor genes. Clusters were identified within GenBank annotated genomes through sequence homology searches and graphical representation using the programs cblaster and clinker, respectively. Homologous genes are represented using a color code and gene links represent an identity threshold of over 30%. The locus accession number is given below the species name.

**Figure S5.**
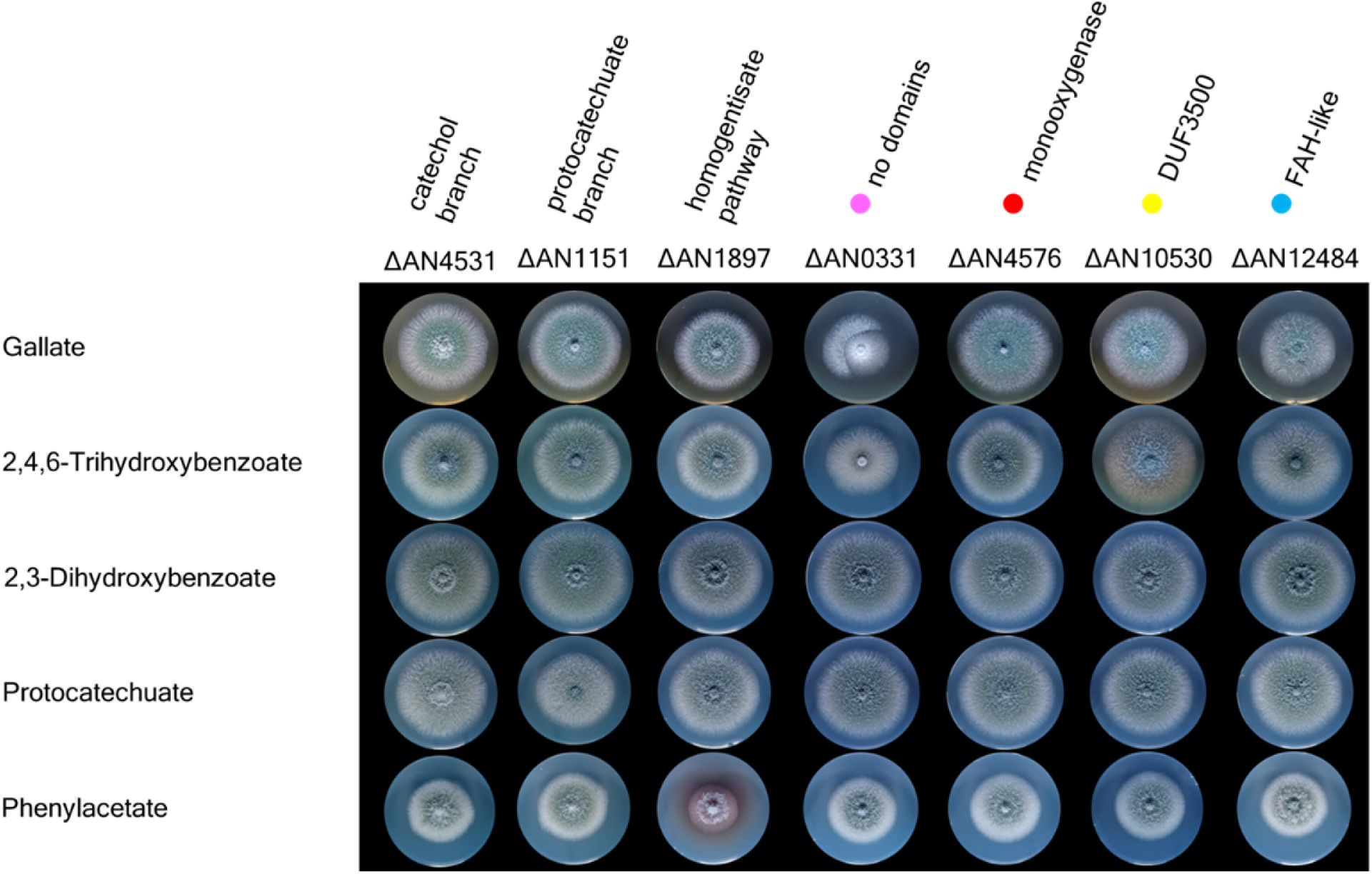
*Aspergillus nidulans* gene deletion mutants grown in gallate and other carbon sources. The four genes of the gallate pathway showed phenotypes of growth impairment for gallate and its isomer 2,4,6-trihydroxybenzoate (except for AN4576) utilization as a carbon source. The mutants used as control of the catechol branch or protocatechuate branch of the 3-oxoadipate pathway, and homogentisate pathway showed the expected growth phenotypes in the respective metabolites of their pathways, *i.e.*, normal growth for ΔAN4531 in 2,3-dihydroxybenzoate, growth impairment for ΔAN1151 in protocatechuate and growth impairment and accumulation of a colored metabolite for ΔAN1897 in phenylacetate. Solid minimal media supplemented with aromatic hydrocarbons (10 mM) and 0.2% (m/v) yeast extract and 0.1% (m/v) casamino acids (pH 6.0).

**Figure S6.**
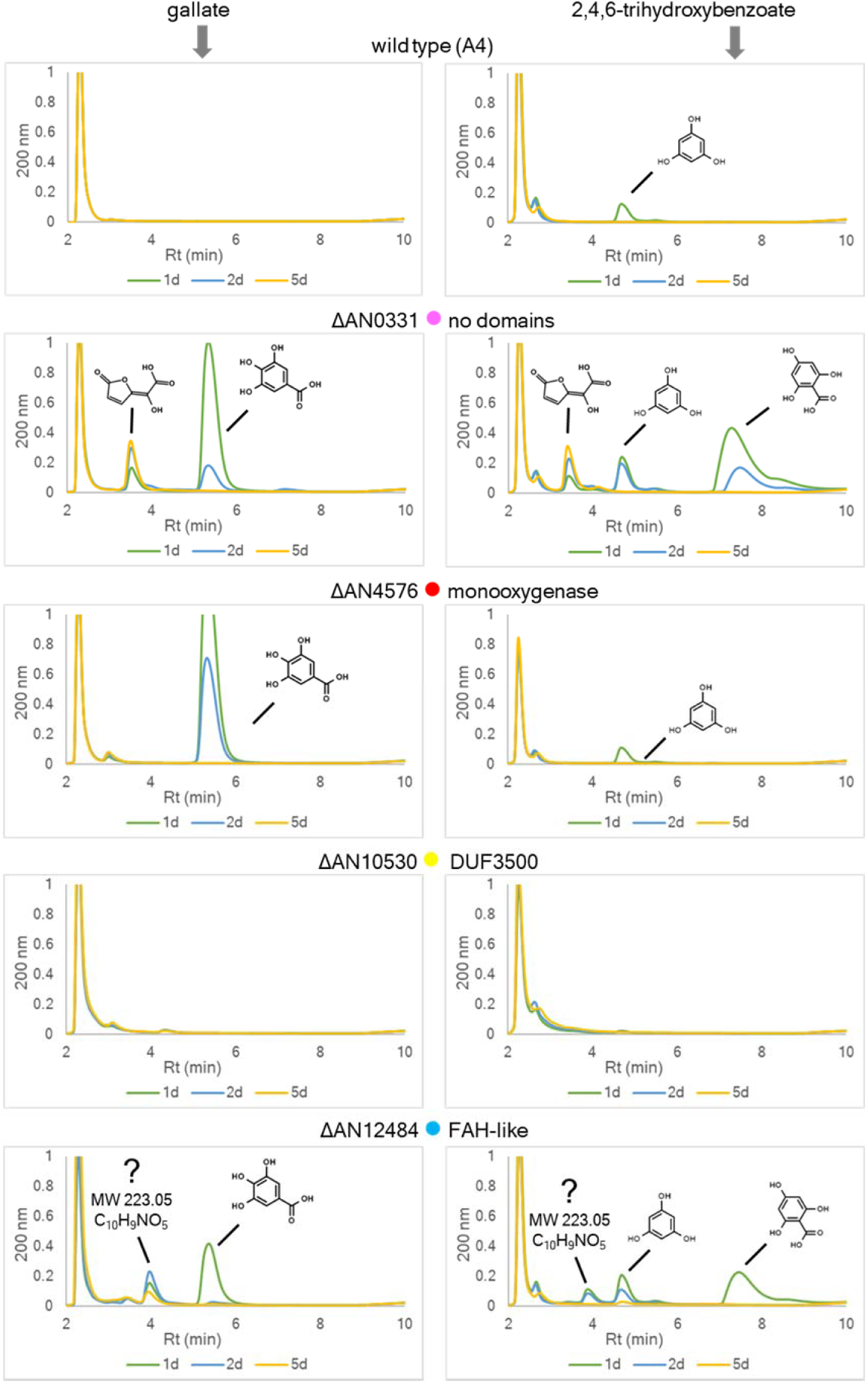
Metabolites differentially accumulated in cultures of the four catabolic genes mutants in the presence of gallate (**left side**) and its isomer 2,4,6-trihydroxybenzoate (**right side**). Note: HPLC method A.

**Figure S7.**
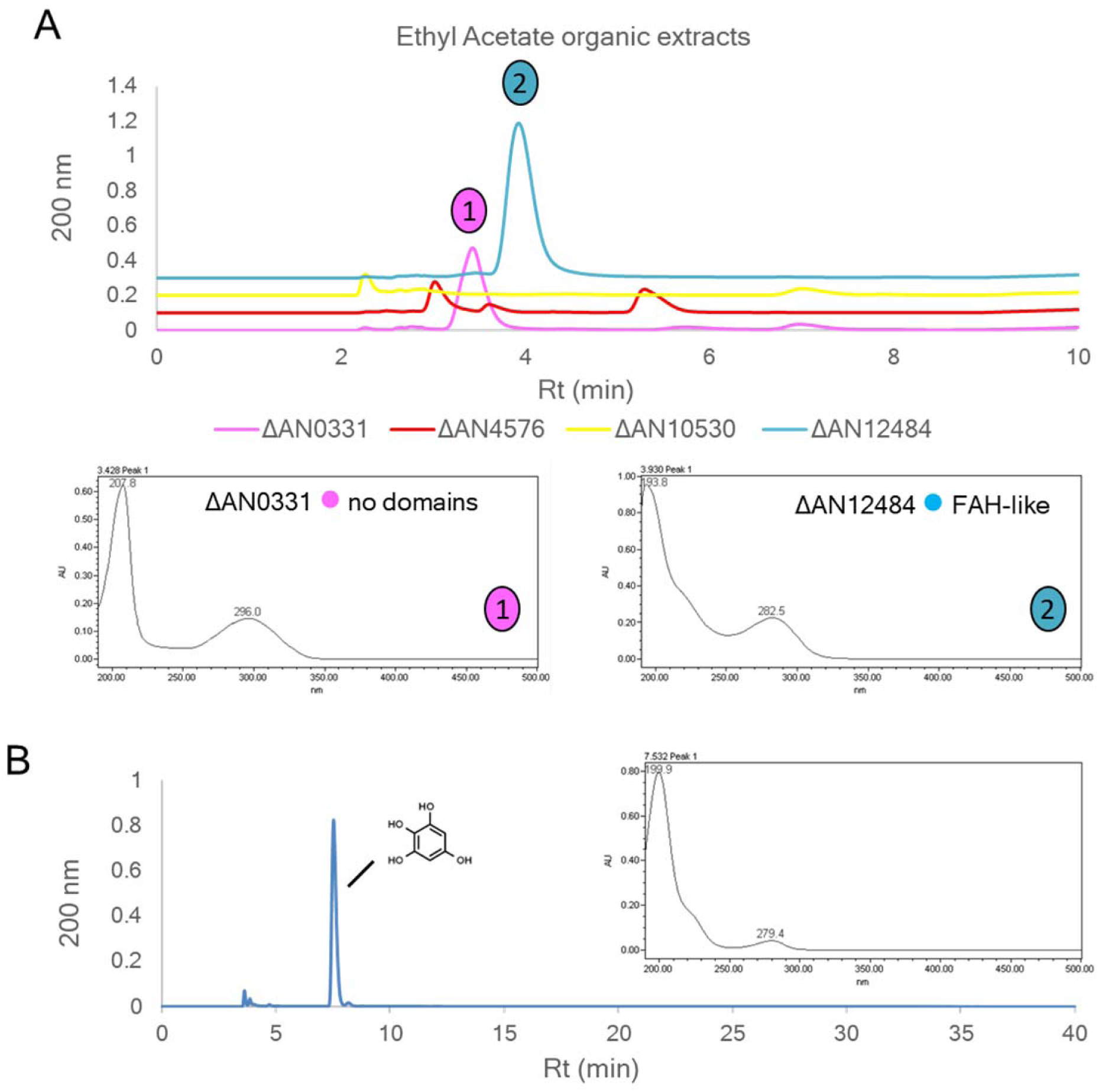
Chromatographic profiles of ethyl acetate extracts (50x) derived from cultures of the four catabolic gene mutants in the presence of gallate (**panel A**; HPLC method A) and of 1,2,3,5-tetrahydroxybenzene (**panel B**; HPLC method B). The absorption spectrum is shown for selected metabolites.

**Figure S8.**
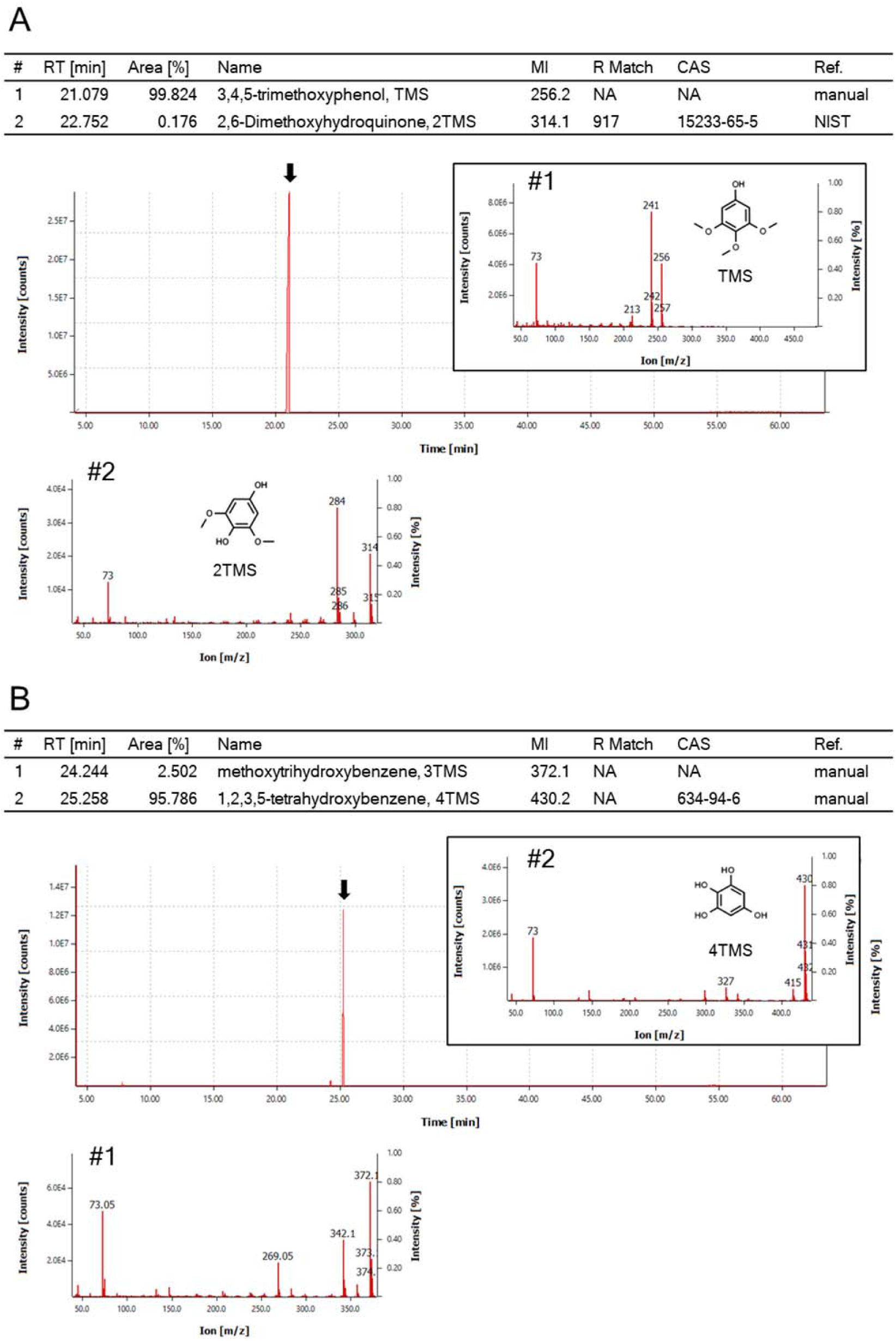
GC-MS chromatograms and the mass spectrum are shown for analyte molecules monitored in the synthesis of 1,2,3,5–tetrahydroxybenzene (panel B) from 3,4,5– trimethoxyphenol (panel A). Analyte molecules exceeding 2% of the chromatogram area or identified with high probability were selected. Abbreviations and notes: retention time (RT); molecular ion (MI), an asterisk indicates as putative; Reverse Match Factor (R Match); reference (Ref.); Chemical Abstracts Service (CAS) Registry Number of the underivatized parent compound; not available (NA); National Institute of Standards and Technology (NIST) Mass Spectral Library.

**Figure S9.**
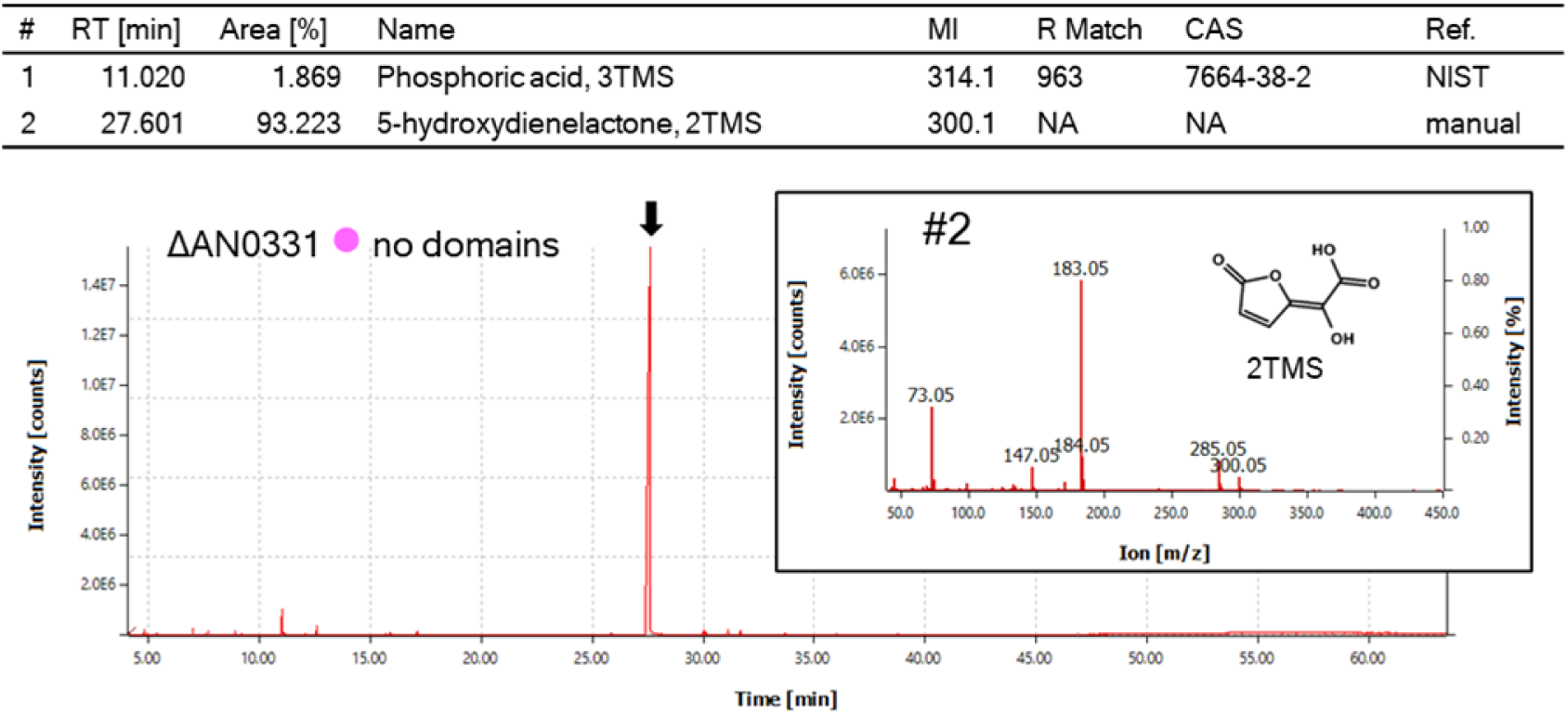
Gallate-derived metabolites differentially accumulated in cultures of the mutant ΔAN0331 (short no domains protein gene). GC-MS chromatograms of the mutant’s culture extract and the mass spectrum of selected analyte molecules are shown. Analyte molecules exceeding 2% of the chromatogram area or identified with high probability were selected. Abbreviations and notes: retention time (RT); molecular ion (MI), an asterisk indicates as putative; Reverse Match Factor (R Match); reference (Ref.); Chemical Abstracts Service (CAS) Registry Number of the underivatized parent compound; not available (NA); National Institute of Standards and Technology (NIST) Mass Spectral Library.

**Figure S10.**
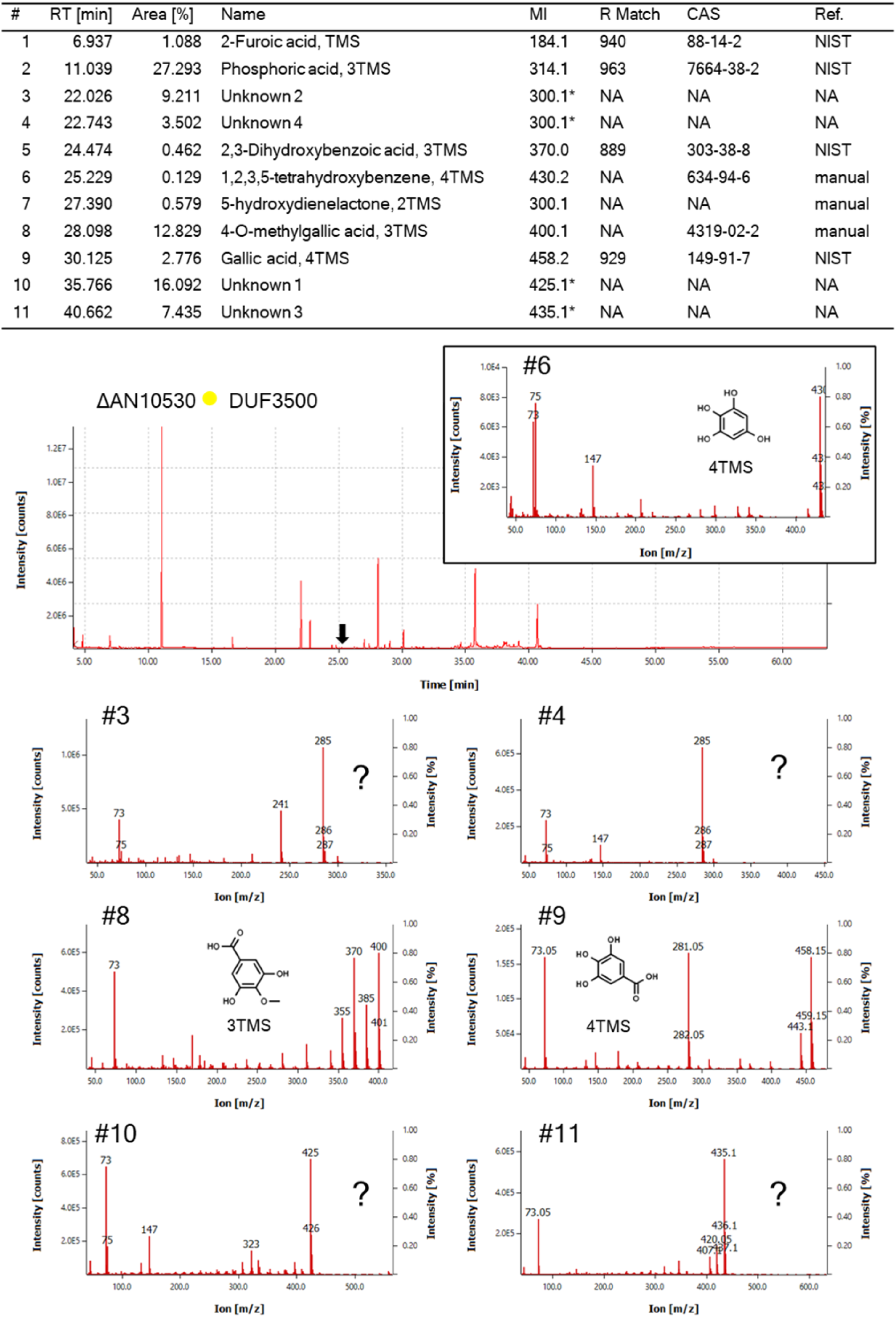
Gallate-derived metabolites differentially accumulated in cultures of the mutant ΔAN10530 (DUF3500 gene). GC-MS chromatograms of the mutant’s culture extract and the mass spectrum of selected analyte molecules are shown. Analyte molecules exceeding 2% of the chromatogram area or identified with high probability were selected. Abbreviations and notes: retention time (RT); molecular ion (MI), an asterisk indicates as putative; Reverse Match Factor (R Match); reference (Ref.); Chemical Abstracts Service (CAS) Registry Number of the underivatized parent compound; not available (NA); National Institute of Standards and Technology (NIST) Mass Spectral Library.

**Figure S11.**
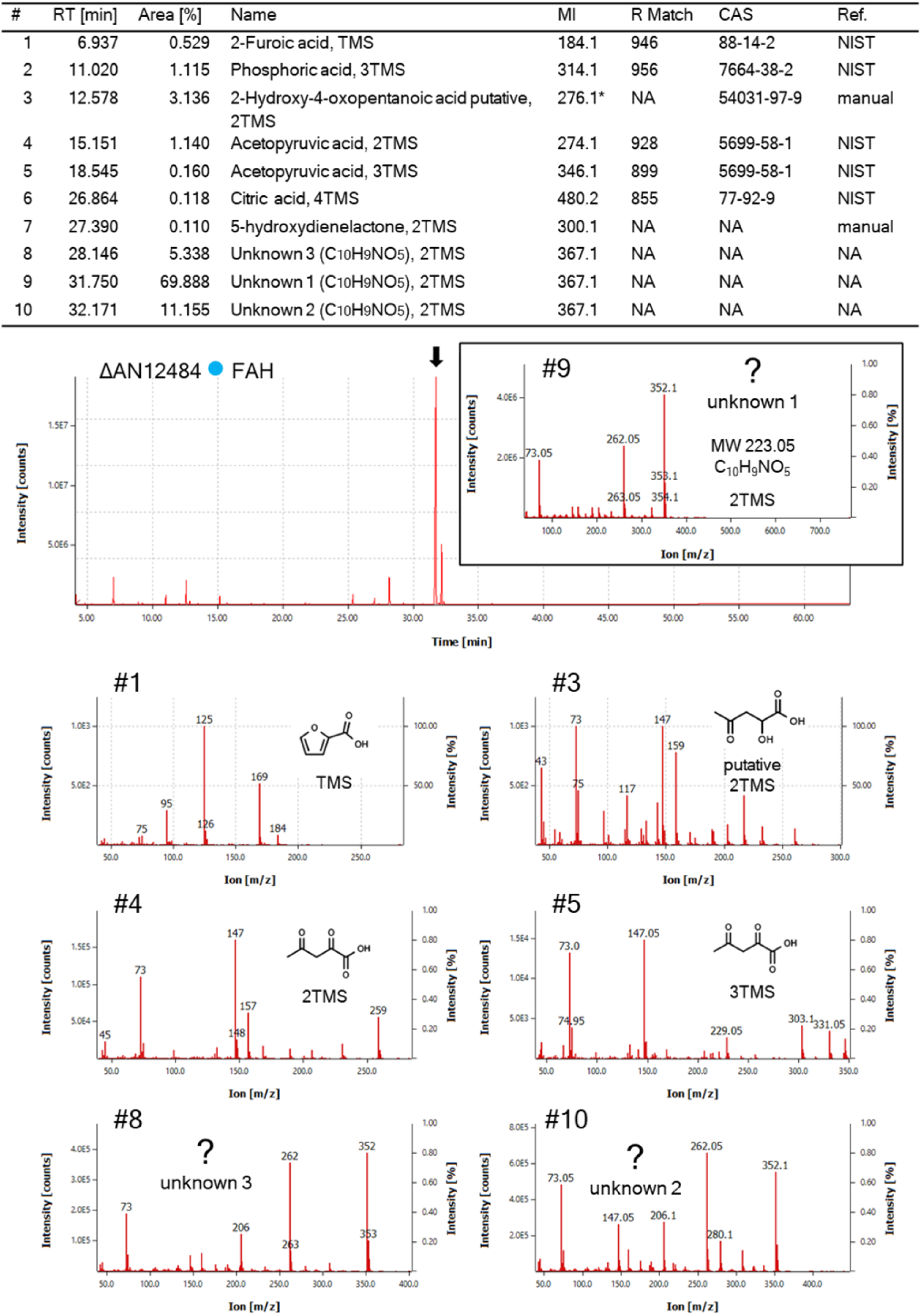
GC-MS analysis of gallate-derived metabolites differentially accumulated in the cultures of the mutant ΔAN12484 (FAH-*like*). GC-MS chromatograms of the mutant’s culture extract and the mass spectrum of selected analyte molecules are shown. Analyte molecules exceeding 2% of the chromatogram area or identified with high probability were selected. Abbreviations and notes: retention time (RT); molecular ion (MI), an asterisk indicates as putative; Reverse Match Factor (R Match); reference (Ref.); Chemical Abstracts Service (CAS) Registry Number of the underivatized parent compound; not available (NA); National Institute of Standards and Technology (NIST) Mass Spectral Library.

**Figure S12.**
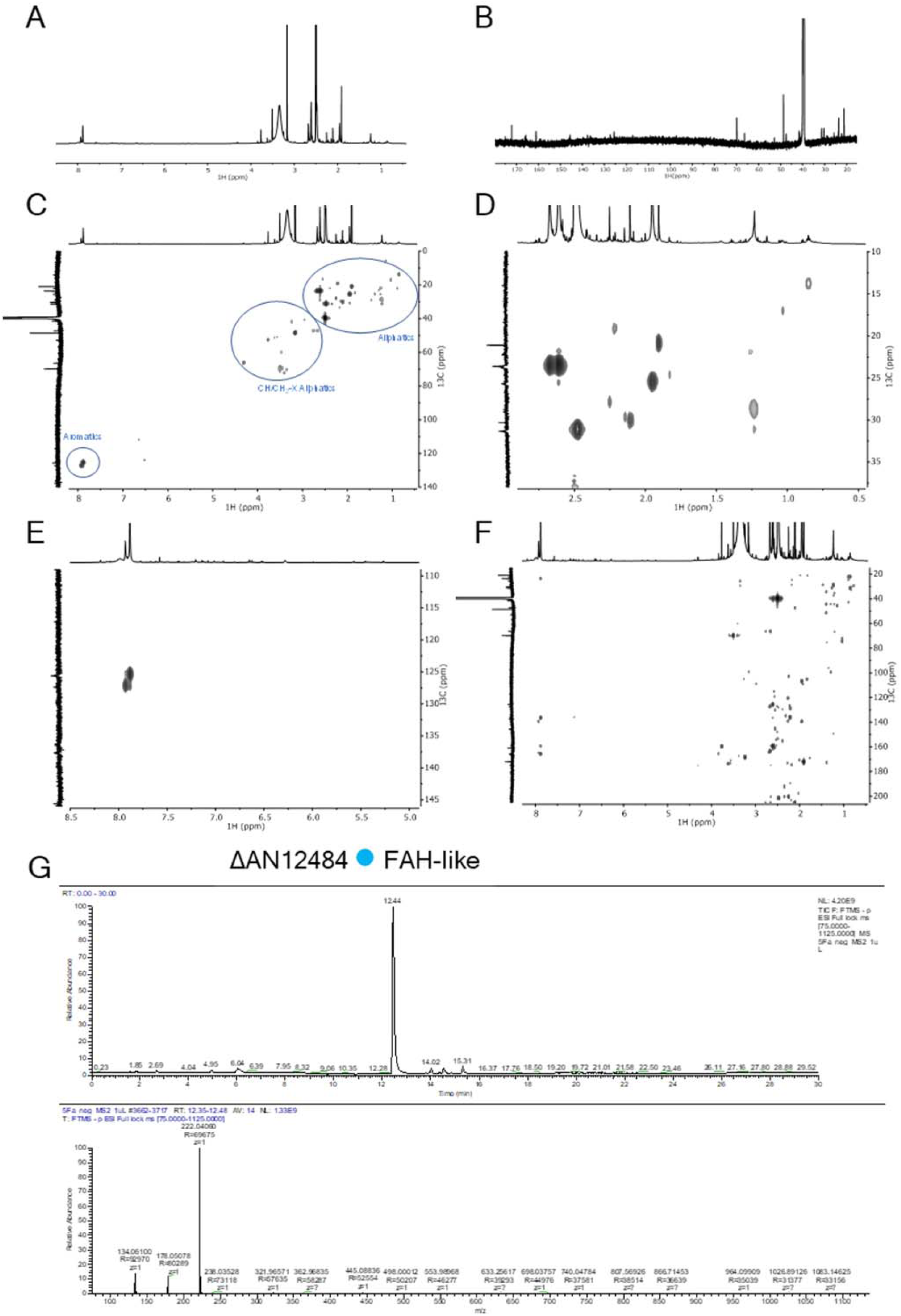
NMR and LC-MS analysis of gallate-derived metabolites differentially accumulated in cultures of the mutant ΔAN12484 (FAH-*like*). Wide-ranging NMR spectral characterization of ΔAN12484 culture extract (**panel A to F**). The ^1^H NMR (A); The ^13^C NMR (B); the ^1^H-^13^C HSQC spectrum (C); HSQC regions corresponding to aliphatics (D) and aromatics (E) and the ^1^H-^13^C HMBC spectrum (F). LC-MS TIC chromatogram and mass spectrum of selected analyte molecules are shown (**panel G**).

